# The Adaptor Protein 2 (AP2) complex modulates habituation and behavioral selection across multiple pathways and time windows

**DOI:** 10.1101/2022.05.20.492863

**Authors:** Rodrigo Zúñiga Mouret, Jordyn P. Greenbaum, Hannah M. Doll, Eliza M. Brody, Emma L. Iacobucci, Nicholas C. Roland, Roy C. Simamora, Ivan Ruiz, Rory Seymour, Leanne Ludwick, Antonia H. Groneberg, João C. Marques, Alexandre Laborde, Gokul Rajan, Filippo Del Bene, Michael B. Orger, Roshan A. Jain

## Abstract

Animals constantly perceive and integrate information across sensory modalities, and their nervous systems must select behavioral responses appropriate to the current situation and prior experience. Genetic factors supporting this behavioral flexibility are often disrupted in neuropsychiatric conditions, and our previous work revealed the disease-associated *ap2s1* gene critically supports habituation learning in acoustically-evoked escape behavior of zebrafish. *ap2s1* encodes a subunit of the AP2 endocytosis adaptor complex and has been linked to autism spectrum disorder, though its mechanism and direct behavioral importance have not been established. Here, we show that multiple subunits of the AP2 complex regulate acoustically-evoked behavior selection and habituation learning. Furthermore, *ap2s1* biases the choice between distinct escape behaviors in sensory modality-specific manners, and more broadly regulates action selection across different sensory contexts. Using tissue-specific and inducible transgenic rescue, we demonstrate that the AP2 complex functions acutely and in the nervous system to modulate acoustically-evoked habituation learning, suggesting several spatially and/or temporally distinct mechanisms through which AP2 regulates different aspects of escape behavior selection and performance. Altogether, we demonstrate that the AP2 complex coordinates action selection across stimulus modalities and contexts, providing a new vertebrate model for the role of *ap2s1* in human conditions including autism spectrum disorder.

**SIGNIFICANCE STATEMENT:** The *AP2S1* gene has been linked to learning disabilities and autism spectrum disorders (ASD), though the mechanisms underlying its impact on human behavior are unknown. We explored how, when, and where this gene regulates vertebrate behavior, developing a zebrafish model to identify the roles and mechanisms through which *ap2s1* modulates behavior. We find that *ap2s1* regulates simple acoustically-evoked learning, as well as how individuals bias behavioral choice in a wide variety of contexts. We show that *ap2s1* acts at multiple distinct time periods and locations both within and outside of neuronal tissues, revealing the diverse mechanisms and pathways through which it modulates vertebrate behavior.

## INTRODUCTION

Animals must constantly integrate a variety of sensory stimuli to select the behavior best suited for survival. When threatening stimuli such as a looming shadow or an abrupt loud noise are perceived, genetically-encoded mechanisms are essential for assessing the situation and initiating an appropriate response ^1–3^. This decision-making process allows individuals to select the most appropriate behavior from their repertoire depending on the current environmental context, internal state, and prior learning ^2,4–6^. Simple non-associative habituation learning provides experience-dependent flexibility in behavior by allowing animals to shift or reduce their response behavior following repeated inconsequential stimuli ^7^. Deficits in decision-making and habituation learning are often associated with autism spectrum disorders (ASD), attention deficit hyperactivity disorder (ADHD), anxiety disorders, and schizophrenia ^2,8–10^. These conditions all have complex genetic underpinnings, so understanding the genetic mechanisms modulating behavior selection is critical to understand functional neurodiversity ^11–14^.

Many clathrin interactome components have been connected to neuropsychiatric conditions including schizophrenia, bipolar disorder, intellectual disability, and ASD, suggesting that clathrin-dependent processes play central roles in complex behavior regulation ^15–17^. The Adapter Protein 2 (AP2) complex targets specific membrane-associated proteins for clathrin-mediated endocytosis via its α, β, σ, and µ subunits ^18^, and these subunits have been linked to diverse aspects of clinically-relevant behavioral control ^17, 19–21^. Expression levels of the AP2β subunit (*AP2B1*) are altered in the brains of individuals with schizophrenia and major depressive disorder, and AP2 subunit expression is reduced in the brains of untreated individuals with bipolar disorder ^19, 22–24^. Mutations disrupting the AP2µ subunit (*AP2M1*) are associated with severe disruption in cognition and behavior in humans ^21^, while mutations in the AP2σ subunit (*AP2S1*) are associated with behavior changes characteristic of autism spectrum disorders, attention hyperactivity disorder, learning disorders, and cognitive dysfunction ^20, 25–28^. We previously found that the zebrafish AP2σ homolog (*ap2s1*) regulates acoustically-evoked behavior selection and habituation learning, indicating the conserved importance of this gene in modulating behavior across vertebrates ^29, 30^. AP2 is also behaviorally important for invertebrates, where glial AP2σ knockdown in *D. melanogaste*r disrupts circadian behavior ^31^ and mutations disrupting AP2α result in locomotor and coordination defects ^32^. In *C. elegans,* mutations in the *AP2S1* ortholog also impair habituation and enhance sensitivity to mechanosensory stimuli ^33^. Overall, the AP2 complex plays common conserved roles in learning and action selection across species.

The AP2 complex impacts a broad array of targets and cellular processes, providing several potential mechanisms to affect behavior, for instance through developmental circuit assembly, acute neuronal function, or a combination thereof. The AP2 complex has been implicated in neurodevelopment through its regulation of cell polarity during differentiation and development ^34^, developmental Wnt signaling ^35^, cell migration ^36, 37^, neuronal survival ^38^, axonal growth cone outgrowth and guidance ^39^, and dendrite formation and extension ^38, 40^. The four AP2 subunits recognize different sets of targets and cofactors, and multiple additional vertebrate clathrin adaptor complexes allow for AP2 subunit-specific functions during these diverse processes ^18^. Additionally, AP2 activity can regulate acute neuronal function presynaptically through synaptic vesicle recycling ^41–44^ and postsynaptically by regulating internalization, recycling, and localization of neurotransmitter receptors including GABAA receptors and ionotropic glutamate receptors in vertebrates and invertebrates ^45–48^. AP2 subunits are broadly expressed in neurons, glia, and several endocrine tissues, allowing autonomous and non-autonomous nervous system regulation ^31, 49–51^. Thus, the locations and degrees to which AP2 regulates behavior developmentally in constructing neural pathways critical for behavioral selection processes, or acutely by regulating synaptic function, remain uncertain.

Here, we use a zebrafish model to determine AP2 complex requirements for behavioral selection and plasticity, leveraging their performance of discrete motor patterns selected from a relatively simple bout repertoire ^52, 53^. We use a set of AP2 mutants to probe escape behavior selection and habituation, showing multiple subunits of the AP2 complex are required to regulate acoustically evoked escapes. Moreover, we show that *ap2s1* regulates behavior selection across diverse visual, vibrational, and spontaneous contexts, indicating a pervasive impact of the AP2 complex on behavior flexibility. Finally, through tissue-and temporally-specific expression experiments we demonstrate that *ap2s1* acts in neurons to modulate acoustically evoked habituation through an acute mechanism.

## RESULTS

### *ap2s1* modulates habituation, sensitivity, behavioral selection, and behavioral performance of acoustically-evoked escape behaviors

We previously identified *ap2s1* as a critical gene in modulating zebrafish acoustically-evoked escape behavior and habituation ^29^. The original allele identified from this genetic screen (*ap2s1^p1^*^72^, formerly *ignorance is bliss^p172^* ^30^) disrupts the splice donor site of the third intron, producing an in-frame deletion removing amino acids 52-89 of the AP2σ protein (**Fig 1A**). We generated two additional *ap2s1* alleles using the CRISPR/Cas9 system (*ap2s1^hv1^* and *ap2s1^p199^*, **Fig 1A**) to determine if *ap2s1^p172^* retains some function and if other portions of the protein are also critical for regulating acoustically-evoked behavior. The *ap2s1^hv1^* allele deletes amino acids 17 & 18 of the protein while remaining in frame, creating a disruption in close proximity to the critical Arg15 target binding residue ^54^. The *ap2s1^p199^* allele contains a frameshift mutation after amino acid 16 that leads to an early stop codon and a severe truncation of nearly 90% of the normal AP2σ protein, likely rendering it functionally null (**Fig. 1A**) ^30^.

**Figure 1:**
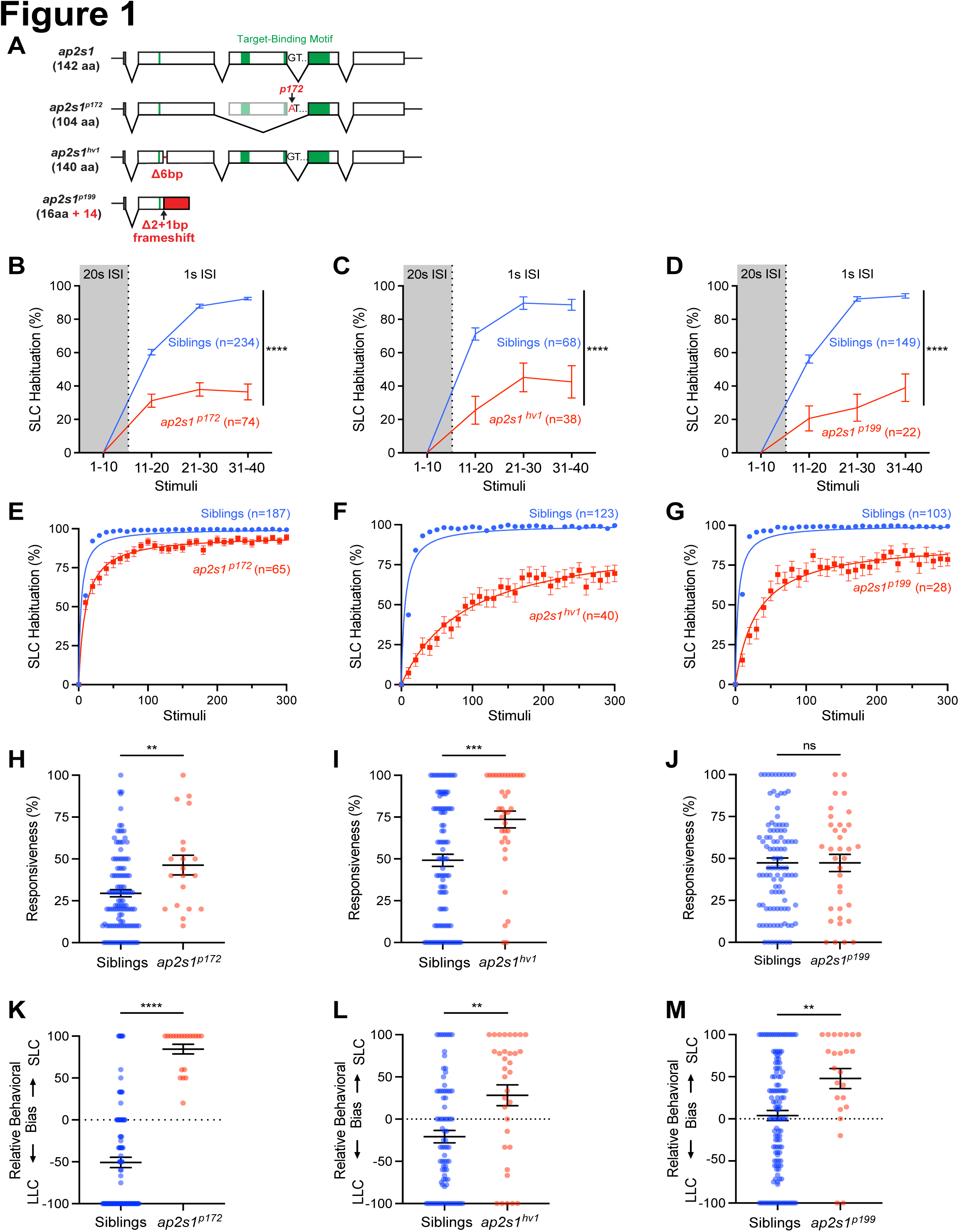
Diverse *ap2s1* alleles disrupt acoustically-evoked habituation learning, responsiveness, and behavior selection bias. **(A)** Diagram of *ap2s1* transcript and alleles, regions contributing to target binding indicated in green with disruptions indicated in red. **(B-G)** Acoustic SLC habituation of sibling (blue) and *ap2s1* mutants (red), presented with 10 intense (23.8 dB) stimuli at 20s ISI, followed by 30 more identical stimuli at 1s ISI. Individual habituation scores were normalized to their baseline SLC response rate at 20s ISI (grey), and *ap2s1* mutants were compared to siblings with a 2-way repeated measures ANOVA (Genotype: ****p<0.0001, Time: ****p<0.0001, Individual: ****p<0.0001). Average SLC habituation scores across 300 identical stimuli at 1s ISI with hyperbolic fitted curves (E-G). **(H-J)** Average responsiveness of individual larvae to 10 low intensity (-0.6 dB) acoustic stimuli presented at 20 s ISI, compared with 2-tailed Mann-Whitney test. **(K-M)** Relative behavioral bias of individual larvae to 10 low intensity (-0.6 dB) acoustic stimuli presented at 20 s ISI, compared with 2-tailed Mann-Whitney test. Mutants were always compared to their corresponding wild type and heterozygous siblings, using *ap2s1^p172^* (B, E, H, K), *ap2s1^hv1^* (C, F, I, L), and *ap2s1^p199^* (D, G, J, M) alleles. All error bars indicate SEM. *p<0.05, **p<0.01, ***p<0.001, ****p<0.0001.

To test whether the new *ap2s1* alleles also disrupted acoustically-evoked habituation learning, we exposed larvae to 10 intense stimuli at 20s interstimulus intervals (ISI) to establish their baseline Short Latency C-Start (SLC) escape response rate, followed by a series of 30 equally intense stimuli at 1s ISI during which SLC escape behavior robustly habituates in wild type animals ^55^. Compared to their siblings, *ap2s1^p172^* mutants showed reduced habituation across the 1s ISI stimuli (**Fig 1B**), consistent with previous findings ^29^. Similarly, both *ap2s1^hv1^* (**Fig 1C**) and *ap2s1^p199^* (**Fig 1D**) significantly impaired acoustic habituation in homozygous mutants, indicating that all three alleles disrupt the ability of AP2σ to regulate acoustic habituation learning. To determine if these habituation defects reflect a reduced rate of habituation and/or a decreased total capacity for habituation, we extended the habituation assay to include 300 identical intense stimuli at 1s ISI, calculated the degree of habituation for each individual in blocks of 10 stimuli across the assay, then fit their habituation to a hyperbolic curve (**Fig. 1E-G**). To assess habituation rate of the mutants, we compared the number of stimuli required to reach half-maximal habituation from the fitted habituation curves. Each of the 3 alleles produced significant increases in the number of stimuli required to reach half maximal habituation (*ap2s1^p172^*: 10.0 vs 3.5 for siblings, p<0.0001; *ap2s1^hv1^*: 90.1 vs 5.3 for siblings, p<0.0001; *ap2s1^p199^*: 31.1 vs 3.9 for siblings, p<0.0001), indicating that all 3 *ap2s1* lesions reduced acoustic habituation rate. To test total habituation capacity, we compared the maximal habituation asymptotes for sibling and mutant habituation curves. While siblings quickly plateaued to 100% habituation in each case, all three *ap2s1* mutations impaired the projected maximal level of habituation reached, indicating that *ap2s1* also regulates the total acoustic habituation capacity (*ap2s1^p172^*: 95.8%, p<0.0001; *ap2s1^hv1^*: 93.4%, p<0.0001; *ap2s1^p199^*: 90.2%, p<0.0001).

We next examined the acoustic responsiveness and acoustically-evoked behavior selection bias of each *ap2s1* allele to determine which lesions disrupt these behaviors, independent of habituation learning. We analyzed responses of homozygous *ap2s1* mutant larvae to weak acoustic stimuli, spaced at 20s ISI to avoid habituation, and compared their response rates to their wild type and heterozygous siblings (**Fig 1H-J**). Both *ap2s1^p172^* and *ap2s1^hv1^* mutants were significantly hyper-responsive to weak stimuli compared to their siblings (p=0.0068 for *ap2s1^p172^*, p=0.0004 for *ap2s1^hv1^*, 2-tailed Mann-Whitney test, **Fig 1H-I**), while we did not observe any responsiveness change in *ap2s1^p199^* mutants relative to their siblings (p=0.8642, 2-tailed Mann-Whitney test, **Fig 1J**). We then examined the acoustically-evoked behavior selection biases of all three *ap2s1* alleles. Larvae select between 2 major escape responses to acoustic stimuli, the SLC and a Long-Latency C-start (LLC), reflecting the assessed threat level of the stimulus and recent stimulus history ^30,56^. All three *ap2s1* alleles shifted the mutant response bias towards SLCs for all stimuli tested (p<0.0001 for *ap2s1^p172^*, p=0.0010 for *ap2s1^hv1^*, p=0.0027 for *ap2s1^p199^*, 2-tailed Mann-Whitney test, **Fig. 1K-M**), together indicating that each lesion disrupts key functions of *ap2s1* in modulating acoustically-evoked responsiveness, behavior selection, and habituation learning.

To determine if *ap2s1* impacts the kinematic performance of the acoustically-evoked escape behaviors selected, we examined the response latency and several key parameters of the initial C-bend of all responses: initial turn angle, initial turn duration, and maximum angular velocity (**Fig 2**). We observed shortened SLC response latency (**Fig 2A, C, E**), mildly reduced SLC turn angles (**Fig 2G, H, I**), and mildly reduced SLC maximal angular velocity (**Fig 2M, N, O**) in all three *ap2s1* mutants compared to their siblings. In contrast, SLC turn duration was not significantly altered for any allele (**Fig 2J, K, L**). LLC responses were less perturbed in *ap2s1* mutants, though we observed increased LLC turn duration for *ap2s1^p172^* (**Fig 2J**), increased LLC maximal angular velocity for *ap2s1^hv1^* and *ap2s1^p199^* (**Fig 2N-O**), and shortened LLC latency for *ap2s1^hv1^* and *ap2s1^p199^* (**Fig 2D, F**). Together, these results reveal that all three mutations disrupt *ap2s1*’s function in regulating escape response kinematics, and that *ap2s1* regulates SLC and LLC response kinematics in different ways. Importantly, we still observed clear and significant kinematic distinctions between SLC and LLC responses in both siblings and *ap2s1* mutants. This indicates that our behavioral classification is robust to these mild kinematic changes and demonstrates that the observed effects of *ap2s1* on habituation learning and behavioral bias are not through altering response kinematics, but rather due to behavioral selection.

**Figure 2:**
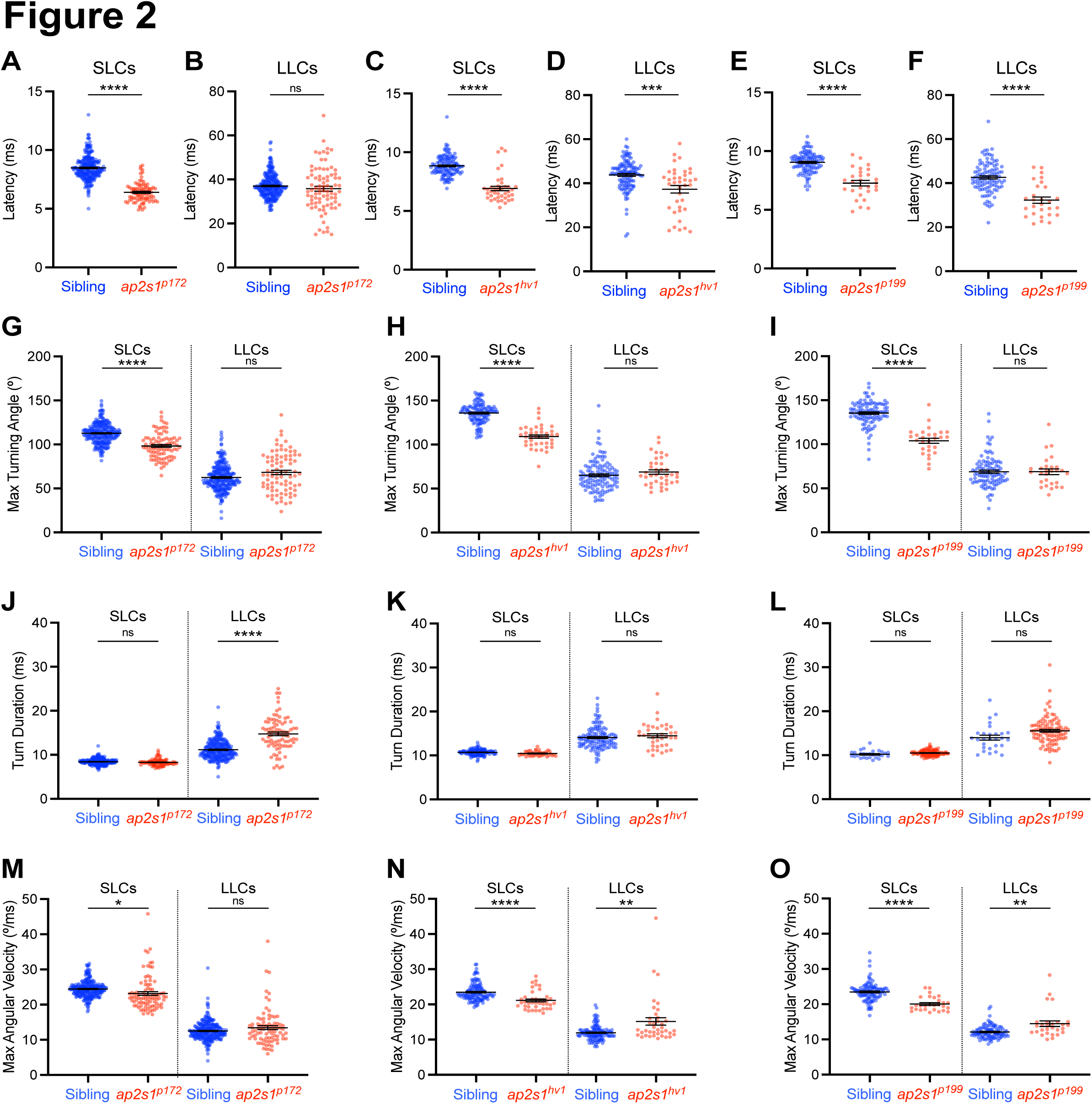
*ap2s1* modulates distinct kinematic features of acoustically-evoked SLC and LLC escape responses. Average movement parameters of acoustically-evoked behavioral responses of sibling (blue) and *ap2s1* mutant (red) larvae across 40 strong (23.8 dB) acoustic stimuli, separated by SLC (left) and LLC (right) responses. **(A-F)** Average acoustic response latencies for SLC (A,C,E) and LLC (B,D,F) behavior of *ap2s1* mutants and their siblings. **(G-I)** Average maximum turning angle of the initial C-bend of *ap2s1* mutant and sibling responses. **(J-L)** Average duration of the initial C-bend of *ap2s1* mutant and sibling responses. **(M-O)** Average maximal angular velocity of SLC and LLC responses of *ap2s1* mutants and siblings. The *ap2s1^p172^* allele was used in panels A, B, G, J, M (n=260 siblings, n=88 mutants), *ap2s1^hv1^* was used in panels C, D, H, K, N (n=123 siblings, n=40 mutants), *ap2s1^p199^* was used in panels E, F, I, L, O (n=100 siblings, n=28 mutants). Error bars indicate SEM. *p<0.05, ***p<0.001, ****p<0.0001, 2-tailed t-test with Welch’s correction.

### Multiple AP2 Complex subunits are critical for acoustically-evoked behavior modulation

Since the four AP-2 subunits can perform different functions within and possibly independent of the full AP2 heterotetromer ^57^, we tested whether *ap2s1*’s impact on acoustic habituation and behavior selection was subunit-specific, or reflective of the function of the entire AP2 complex. We used a point mutation in *ap2a1* that disrupts the splice donor site of the 13^th^ intron, predicted to disrupt the C-terminal Appendage Domain of AP2α, which interacts with many adaptor and accessory proteins during endocytosis (*ap2a1^sa^*^1907^, **Fig 3A**)^58^. Similar to the habituation deficit of *ap2s1* mutants, acoustic habituation was compromised in *ap2a1^sa1907^* mutants compared to their siblings (Genotype: ****p<0.0001, Time: ****p<0.0001, Individual: ****p<0.0001, 2-way repeated measures ANOVA, **Fig 3B**). In response to weak acoustic stimuli, *ap2a1^sa1907^* mutants were both more responsive (p<0.0001, 2-tailed Mann-Whitney test, **Fig 3C**) and more strongly biased toward selecting SLC responses than their siblings (p<0.0001, 2-tailed Mann-Whitney test, **Fig 3D**). To see if *ap2a1* also influences escape response kinematics, we examined the SLC and LLC latencies, turn angles, turn durations, and maximal angular velocities of *ap2a1* mutants vs their siblings. The *ap2a1^sa1907^* mutation resulted in mild kinematic changes to the SLC and LLC behaviors that resembled *ap2s1* disruption (**Fig 3E-I**). Compared to their siblings, *ap2s1^sa1907^* mutants initiated SLC behavior with shortened response latencies (p<0.0001, 2-tailed t-test with Welch’s correction, **Fig 3E**) and a mild decrease in the initial turning angle (p<0.0001, 2-tailed t-test with Welch’s correction, **Fig 3G**), though the initial turn duration (p=0.3031, 2-tailed t-test with Welch’s correction, **Fig 3H**) and maximum angular velocity of SLC responses were not significantly altered (p=0.2472, 2-tailed t-test with Welch’s correction, **Fig 3I**). We also observed a shortening of the initiation latency of LLC behaviors in *ap2s1^sa1907^* mutants (p=0.0002, 2-tailed t-test with Welch’s correction, **Fig 3F**), though the maximum turning angle, turn duration, and maximal angular velocity of LLC behaviors were not significantly impacted in mutants (p=0.9332, p=0.2655, p=0.1971, respectively, 2-tailed t-test with Welch’s correction, **Fig 3G-I**). Nevertheless, SLC and LLC behaviors of *ap2s1^sa1907^* mutants remained kinematically distinguishable from each other, demonstrating that the major acoustically-evoked behavior defects were due to a shift in behavior selection rather than abnormal movement patterns. Taken together, these data show that *ap2a1* modulates acoustic habituation learning and behavior selection bias similarly to *ap2s1*, and thus these likely reflect critical functions of the AP2 complex as a whole.

**Figure 3:**
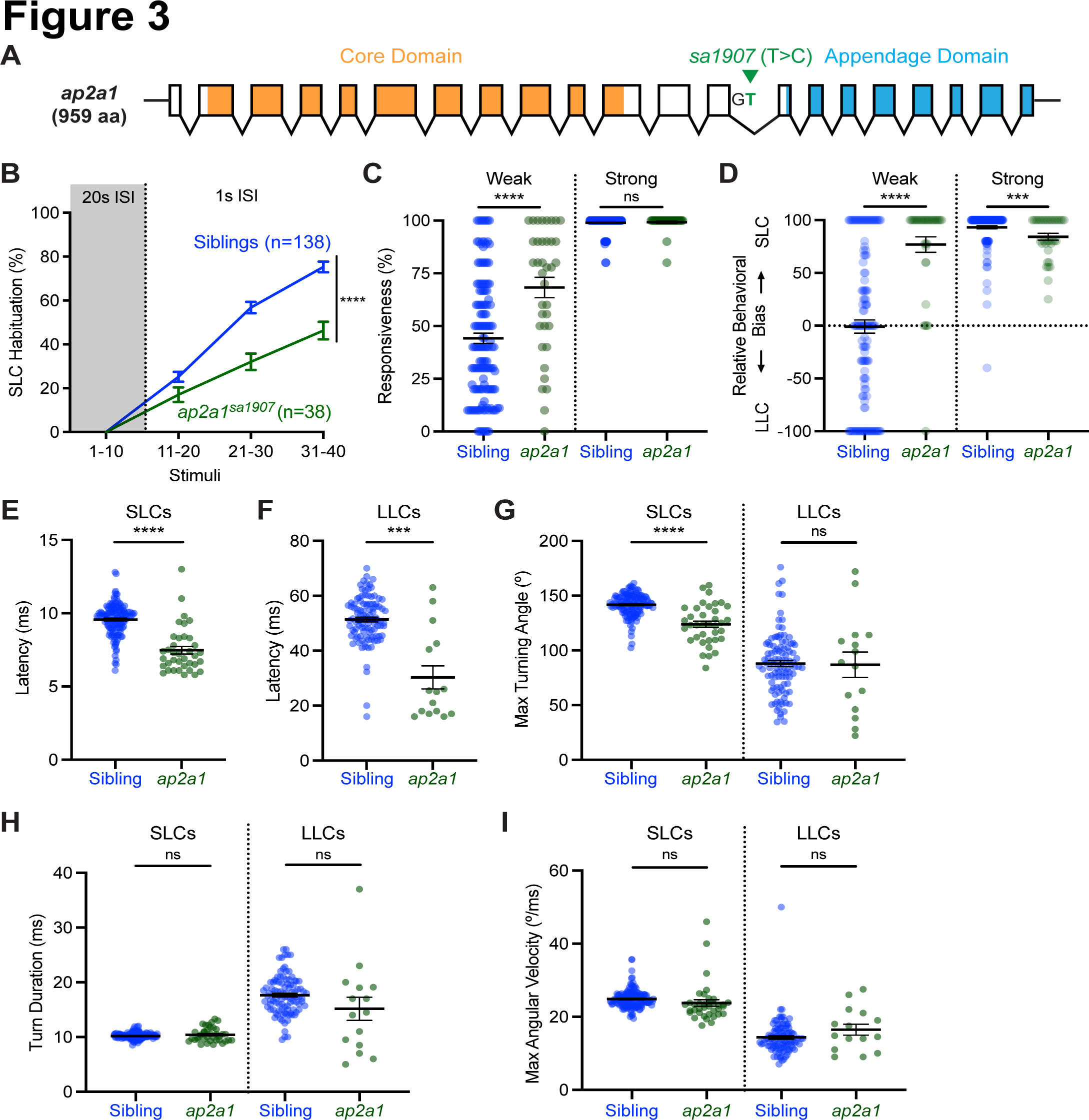
The AP2α subunit regulates acoustically-evoked escape behavior. **(A)** Diagram of *ap2a1* transcript and disrupted splice donor site of *ap2a1^sa1907^* (green). Core domain represented in orange and flexible appendage domain represented in blue. **(B-D)** Acoustically-evoked response modulation of siblings (blue, n=138) and *ap2a1^sa1907^* mutants (green, n=38). Average SLC habituation scores across 30 stimuli at 1s ISI (B), normalized to each individual’s baseline SLC response levels to 10 strong (23.8 dB) stimuli at 20s ISI. *ap2a1^sa1907^* mutants were compared to siblings with a 2-way repeated measures ANOVA (Genotype: ****p<0.0001, Time: ****p<0.0001, Individual: ****p<0.0001). Average responsiveness (C) and relative behavioral bias (D) of individuals to 10 weak (-6.7 dB) and 10 strong (23.8 dB) stimuli at 20 s ISI. ***p<0.001, ****p<0.0001, 2-tailed Mann-Whitney test. **(E-I)** Average kinematic parameters of individual sibling (blue, n=142) and *ap2a1^sa1907^* mutant (green, n=37) larvae in response to 40 strong (23.8 dB) acoustic stimuli, separated by SLC (left) and LLC (right) responses. Parameters measured were the initial response latency (E-F), maximum turning angle of the initial C-bend (G), initial C-bend duration (H), and maximum angular velocity of the initial C-bend (I). ***p<0.001, ****p<0.0001, 2-tailed t-tests with Welch’s correction. All error bars indicate SEM.

### *ap2s1* regulates escape behavior selection in a modality-specific manner

To determine whether *ap2s1* modulates escape behavior selection specifically in acoustic contexts or more broadly across modalities, we examined the escape behavior selection of *ap2s1^p172^* mutants in response to mechanosensory and visual stimuli. First, we exposed larvae to sudden, localized water vibrations which are sensed largely through the lateral line hair cells and can evoke both SLC and LLC escape behaviors (**Fig 4A**) ^59^. Overall, *ap2s1^p172^* mutants were less responsive to these stimuli than their siblings (**Fig 4B**), though they still performed both SLC and LLC behaviors (**Fig 4C-D**). In contrast to their behavioral bias following acoustic stimuli, disrupting *ap2s1* reduced the likelihood of selecting SLC responses (p<0.0001, 2-tailed t-test with Welch’s correction, **Fig 4C**), while overall the rate of LLC response performance was unperturbed (p=0.5424, 2-tailed t-test with Welch’s correction, **Fig 4D**). As a result, the relative behavioral bias of *ap2s1^p172^* mutants showed a significant shift toward LLC escapes in response to local water vibrations (p<0.0001, 2-tailed Mann-Whitney test, **Fig 4E**).

**Figure 4:**
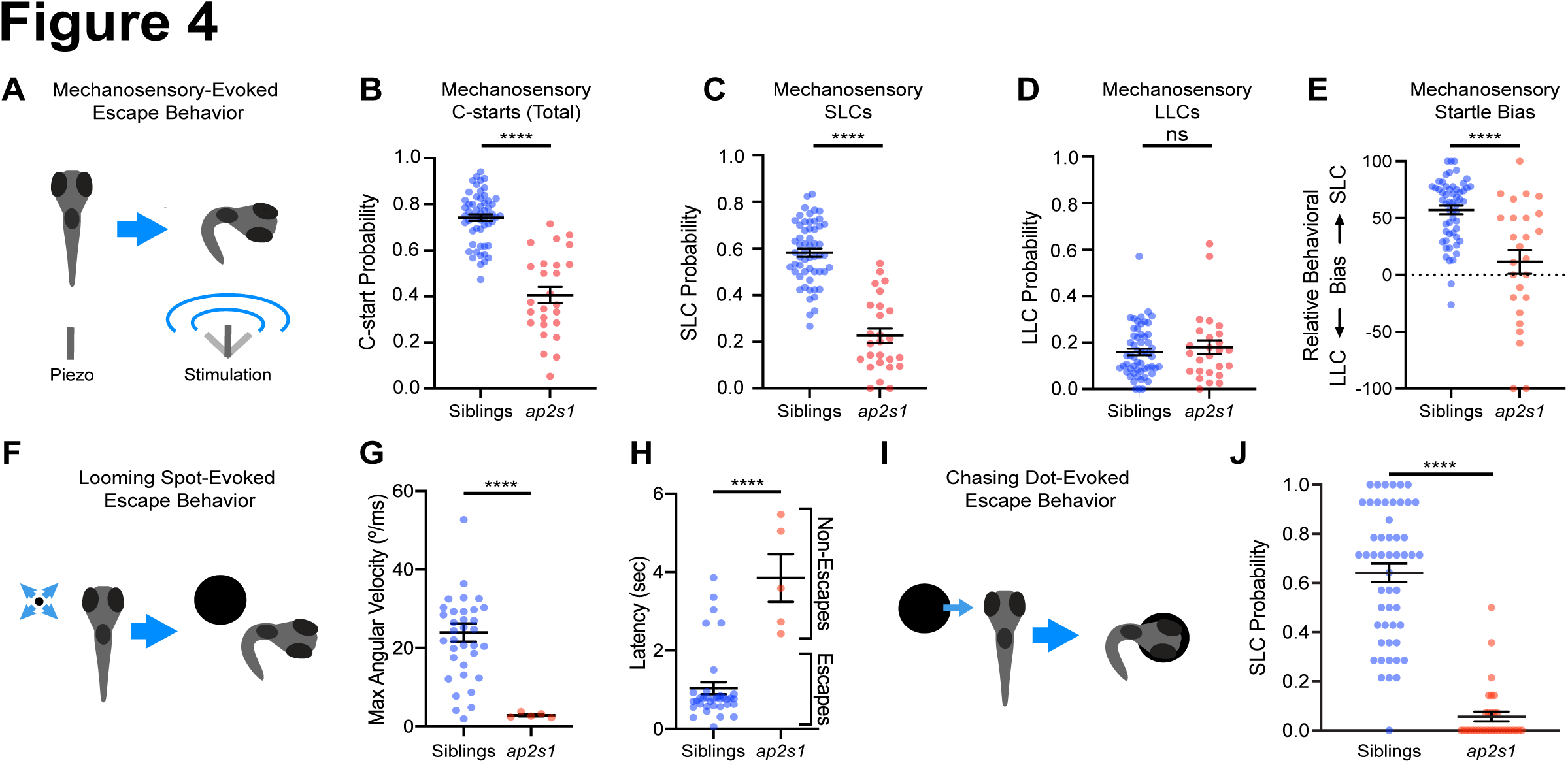
*ap2s1* differentially regulates SLC and LLC escape behavior across varied stimulus modalities. (**A-B**) C-start response probability of *ap2s1^p172^* mutants (red, n=26) to vibrating piezo stimuli compared to non-mutant siblings (blue, n=56). ****p<0.0001, 2-tailed t-test with Welch’s correction. **(C-E)** Within mechanosensory induced C-start responses, the probabilities of SLC responses (C) and LLC responses (D) for siblings (blue) and *ap2s1^p172^* mutants (red), ****p<0.0001, ns = not significant, 2-tailed t-test with Welch’s correction. Relative behavioral bias of mechanosensory-evoked escape behaviors (E) for siblings and *ap2s1^p172^* mutants, ****p<0.0001, Mann-Whitney test. **(F-H)** Maximum angular velocity (G) and response latency (H) of individual behavioral responses of siblings (blue, n=5) and *ap2s1^p172^* mutants (red, n=5) to looming visual stimuli. ****p<0.0001, 2-tailed student’s t-test with Welch’s correction. **(I-J)** Average SLC frequency of non-mutant siblings (blue, n=53) and *ap2s1^p172^* mutants (red, n=33) in response to visual “chasing dot stimuli.” ****p<0.0001, 2-tailed Mann-Whitney test. Error bars represent SEM.

We next examined if *ap2s1* regulated visually-evoked escape behavior selection, using a looming visual stimulus paradigm which, like acoustic stimuli, can evoke both Mauthner-dependent and -independent escape behavior ^60,61^. Larvae were presented with expanding black spots (“looming” visual stimuli), projected from below and locked to the instantaneous position and orientation of the fish throughout the assay (**Fig 4F**). Surprisingly, while these looming stimuli reliably evoked high-velocity short-latency escape responses from siblings, *ap2s1^p172^* mutants failed to perform any high-velocity short-latency escape responses to these stimuli (**Fig 4G-H**). To determine if this lack of any escape responses might simply indicate that *ap2s1* mutant fish are blind to visual stimuli, we tested a second visual context which can evoke escape behavior: a dark spot approaching at constant speed (**Fig 4I**) ^59^. This "chasing dot" stimulus reliably evoked escape behavior in sibling larvae (63.46±3.18%), while overall escape behavior of *ap2s1^p172^* larvae was reduced though not abolished (5.62±1.96%), indicating that visual spot stimuli can indeed be detected by *ap2s1* mutant larvae. Whereas siblings show a strong bias toward responding to the chasing dot stimuli with SLCs, *ap2s1* mutants were much less likely to perform SLC escape behavior to these chasing dot visual stimuli (p<0.0001, 2-tailed Mann-Whitney test, **Fig 4J**). Taken together, these data indicate that *ap2s1* plays a critical role in regulating escape behavior selection in each modality tested, though the direction of its effect on behavior bias is specific to stimulus modality.

### *ap2s1* broadly regulates larval behavior selection across environmental contexts

Given the modality-specific shifts in escape behavior of *ap2s1* mutants, we hypothesized that *ap2s1* may play a broad role in behavioral selection, beyond escape behaviors. Zebrafish larvae select between discrete movement bouts from a simple motor repertoire to navigate their environment ^53,62^. We first examined the role of *ap2s1* in selecting responses to whole-field illumination changes. Larvae typically respond to sudden whole field light reductions (Dark Flashes, DFs) with characteristic large-angle reorientation bouts which are independent of the acoustic escape circuitry, including O-bends and Spot Avoidance Turns (SATs) ^53,63,64^. *ap2s1^p199^* mutant and sibling larvae showed similar response rates to DFs presented at 150s ISI, indicating that mutants can detect this visual stimulus (93.15±1.13% in siblings vs 92.85±2.07% in mutants, p=0.89, 2-tailed Mann-Whitney test, **Fig 5A**). In contrast to the decreased latency of acoustically-evoked startles in *ap2s1* mutants, *ap2s1* mutants initiated DF responses at significantly longer latencies than their siblings **(Fig 5B)**. These mutant response latencies were bimodally distributed (p<0.0001, Hartigan’s dip test for unimodality/multimodality, D=0.114), suggesting that *ap2s1* mutants shifted their DF-evoked responses to kinematically distinct longer-latency turning bouts. To distinguish between these responses, which are more kinematically continuous than acoustically-evoked behavior ^65^, we classified sibling and *ap2s1^p199^* DF responses into 13 kinematically distinct swim and reorientation bouts using a previously described approach ^53^ that we adapted for offline tracking and differences in frame rate and spatial resolution **(Fig 5C)**. The most common turn bouts selected by siblings in response to DF stimuli were SATs (41.1%, **Fig 5D**), LLCs (19.4%, **Fig 5E**), Routine Turns (9.2% RTs, **Fig 5F**) and O-bends (9.0%, **Fig 5G**). In contrast, *ap2s1^p199^* mutants shifted their response profile to select significantly fewer SATs (p<0.0001, 2-tailed Mann-Whitney test, **Fig 5D**) and more RTs at longer latency instead (p<0.0001, 2-tailed Mann-Whitney test, **Fig 5F**), with no significant change in LLC or O-bend response frequency (p=0.3671, p=0.9208, respectively, 2-tailed Mann-Whitney tests, **Fig 5E,G**). To confirm that the observed change in bout selection was not caused by misclassifying mutant movements due to a general shift in movement kinematics, we compared the distributions of bouts in kinematic space of both *ap2s1^p199^* mutants and siblings to previous data used to assign bout categories^53^, finding consistent overlap (**Fig S1A)**. Furthermore, in the same PCA space the *ap2s1^p199^* mutant and sibling distributions showed the specific increase in RTs and decrease in SATs affecting the full kinematic range of each bout type, outside of the region of overlap (**Fig S1B**). Finally, to determine if *ap2s1* regulates visual habituation in this context, we presented the larvae with an additional 42 Dark Flash stimuli at 15s ISI to provoke short-term habituation of the O-bend response, which manifests as a reduction in total response rate to DFs as well as a weakening of kinematic parameters including latency ^55,64^. Since we had observed differences in bout selection between *ap2s1* mutants and siblings following non-habituating stimuli, we grouped all large-angle turns together and examined their habituation to repeated DF stimuli. Compared to their siblings, *ap2s1^p199^* mutants showed enhanced habituation of large angle turns (two-way repeated measures ANOVA, Genotype p=0.0043, Time p<0.0001, Individual p<0.0001, **Fig 5H**) indicating that *ap2s1* may normally act to decrease this aspect of visually-evoked habituation, in contrast to its function increasing acoustically-evoked habituation, and thus *ap2s1*’s impact on short term habituation is modality-specific.

**Figure 5:**
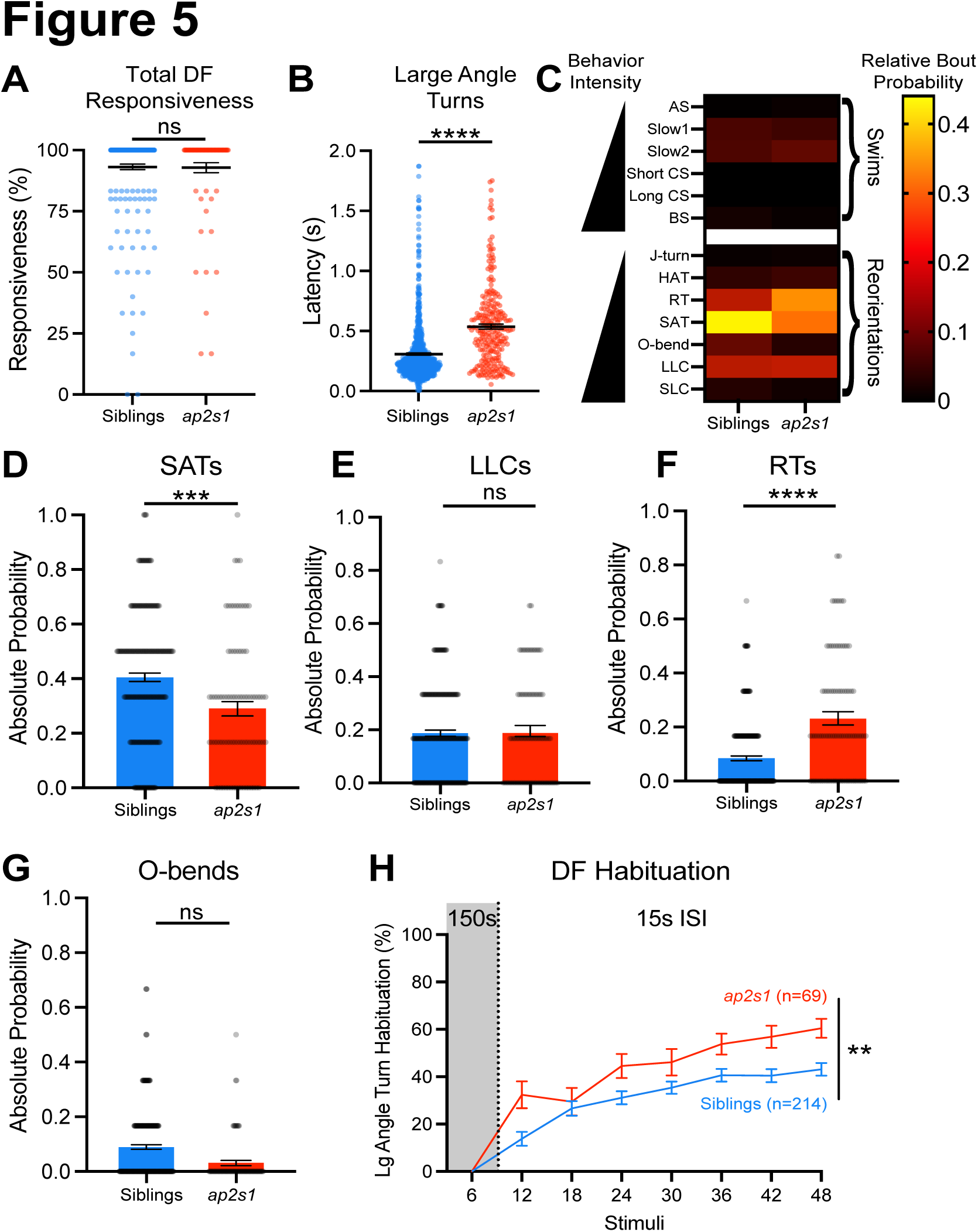
*ap2s1* modulates visually-evoked behavior selection and habituation learning. (**A**) Total responsiveness (A) to 6 dark flash (DF) stimuli presented at 150 s ISI by sibling (blue, n=234) and *ap2s1^p199^* mutant larvae (red, n=79), 2-tailed Mann-Whitney test. Response latency of large angle turns (>95°) by siblings (blue) and *ap2s1^p199^* mutants (red) following DF stimuli in A (2-tailed Mann-Whitney test, ****p<0.0001). (**B**) Relative frequencies of 13 behavioral bout types in response to DFs between siblings (blue) and *ap2s1^p199^* mutants (red). Forward swims and turning bouts are placed roughly in order of increasing vigor, as classified in Marques et al^53^, including Approach Swim (AS), Slow Swim 1 (Slow 1), Slow Swim 2 (Slow 2), Short Capture Swim (SCS), Long Capture Swim (LCS), Burst Swim (BS), J-turn (J-turn), High Angle Turn (HAT), Routine Turn (RT), Spot Avoidance Turn (SAT), O-Bend (O-Bend), Long Latency C-start (LLC), and Short Latency C-start (SLC). **(D-G)** Absolute probabilities of DF response behaviors by siblings (blue) and *ap2s1^p199^* mutants (red), focusing on SATs (D, Mann-Whitney U=7842, 2-tailed Mann-Whitney test, ***p=0.0001), LLCs (E, Mann-Whitney U=10110, 2-tailed Mann-Whitney test, p=0.367), RTs (F, Mann-Whitney U=6565, 2-tailed Mann-Whitney test, ****p<0.0001), and O-bends (G, Mann-Whitney U=570.5, 2-tailed Mann-Whitney test, p=0.921). See also Supplemental Figure S1 for details. (**H**) Habituation of large angle turn responses of siblings (blue, n=214) and *ap2s1^p199^* mutants (red, n=69) to eight blocks of six identical DF stimuli, presented at 150s ISI for the first block and 15s ISI for subsequent blocks. Habituation assessed by two-way repeated measures ANOVA (Genotype **p=0.0043, Time ****p<0.0001, Individual ****p<0.0001). Error bars represent SEM in all panels.

To examine if *ap2s1* regulates more complex behavioral scenarios where larvae must coordinate sequential patterns of movement bouts, we tested if disrupting *ap2s1* impacted the optomotor response (OMR) and hunting behavior. To assess the role of *ap2s1* in optomotor behavior, we tracked mutant and sibling motion when reacting to directionally moving gratings projected below. *ap2s1^p172^* mutants and their non-mutant siblings both move in the correct direction along with the stimuli significantly more than they would swim that direction on average (p<0.0001 and p=0.020, respectively, one sample t-test, **Fig 6A**), indicating that they can both perceive and respond to OMR stimuli. Mutants choose to move in the correct direction more often than siblings during this assay (p=0.0216, Mann-Whitney U=180, two-tailed Mann Whitney test, **Fig 6A-B**). To assess if *ap2s1* influences goal-directed hunting behaviors we examined *ap2s1^p172^* mutants in a rotifer consumption assay, in which larvae must execute a complex sequence of visually-guided behaviors to successfully track and capture prey ^66–68^. While siblings consumed 15.0% of available rotifers during the 135-minute hunting period, *ap2s1^p172^* mutants did not appreciably consume rotifers during this period, indicating deficits in visually-guided hunting ability (p=0.00012, unpaired t-test with Welch’s correction, **Fig 6C**). Notably, we have successfully raised homozygous *ap2s1^p172^* mutants past 90 days old on a rotifer and brine shrimp diet, indicating that *ap2s1* mutants do retain the capacity to hunt and consume prey over longer time periods than this assay. Combined, these data reveal specific abnormalities across a diverse array of behavior selection in simple and complex visually-guided contexts.

**Figure 6:**
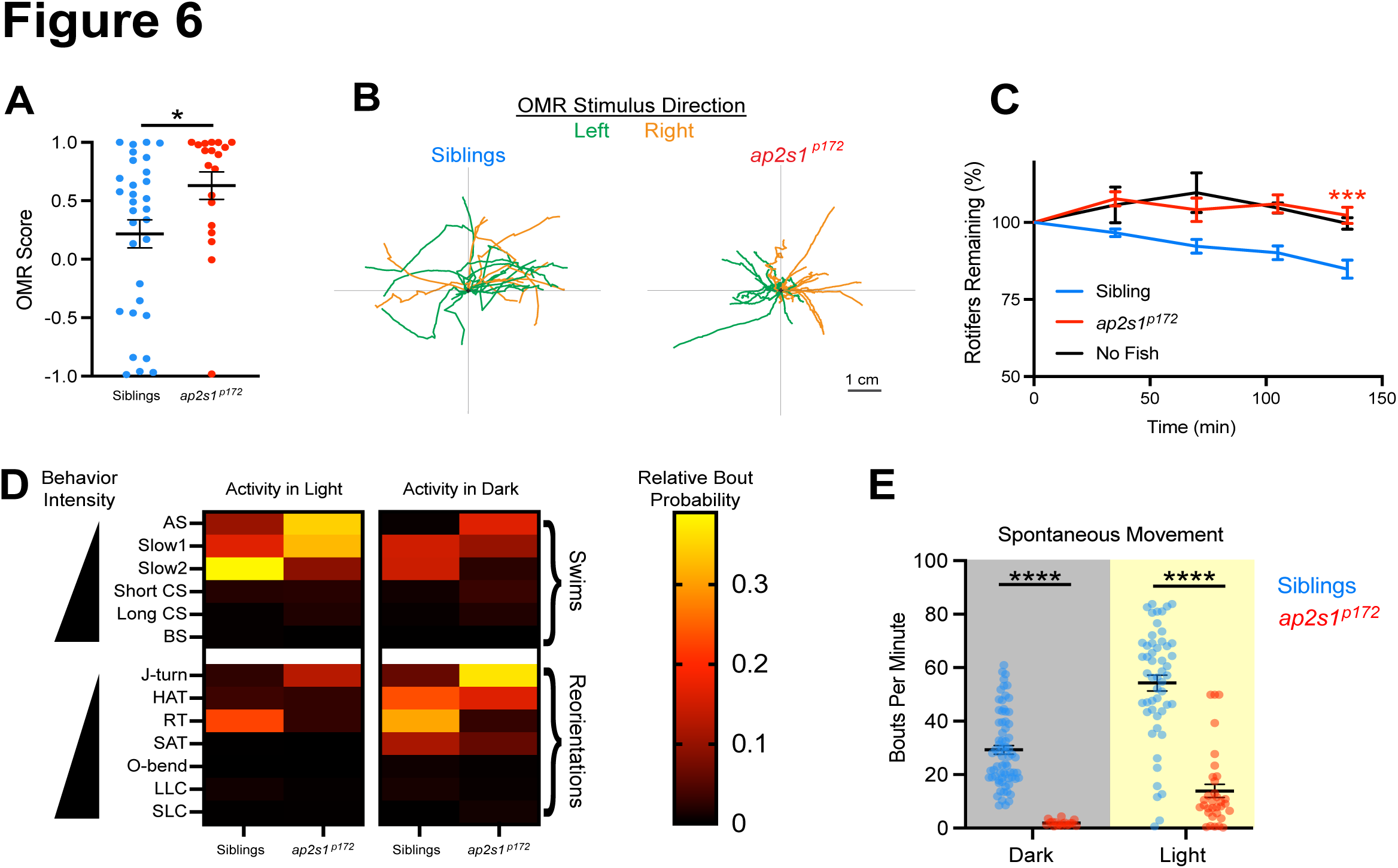
ap2s1 regulates behavioral performance and choice in goal-directed and spontaneous contexts. **(A)** OMR score of siblings (blue, n=16) and *ap2s1^p172^* mutants (red, n=11) to moving optomotor visual stimuli, where positive scores indicate overall movement in the same direction as the visual stimuli. *p<0.05, 2-tailed Mann-Whitney test. **(B)** Relative fish movement trajectories following the onset of each leftward (green) or rightward (orange) optomotor stimulus, depicted starting from the black cross at the center of each graph. Presented paths of siblings (left graph) and mutants (right graph) were truncated upon exiting a square region of interest during the trial (see Materials & Methods). (**C**) Relative consumption of rotifers by non-mutant siblings (blue, n=23), *ap2s1^p172^* mutants (red, n=11), and control arenas with no fish (black, n=3) over a 135-minute time period (***p≤0.001, 2-tailed t-test with Welch’s correction, at conclusion of assay). Rotifer counts for each individual were normalized to 100% at time 0. **(D-E)** Sibling and *ap2s1^p172^* mutant spontaneous behavior in the light (n=52 siblings, n=33 mutants) and the dark (n=72 siblings, n=18 mutants). ****p<0.0001, 2-tailed Mann-Whitney test. See also Supplemental Figure S2 for details. Error bars represent SEM.

Finally, we assessed whether *ap2s1* impacts behavior selection during spontaneous activity in the absence of acute stimuli by recording and classifying the behavioral bout choice of spontaneously active larvae in light and dark environments, following the classification scheme of Marques et al. (2018). In both environments, *ap2s1^p172^* mutants significantly shifted their selection of behavioral bouts, tending to select lower amplitude forward swims and turns (**Fig 6D, Fig S2**). In both illumination contexts, *ap2s1^p172^* mutants showed a significant enrichment for J-turn behavior (dark: p<0.000001, light: p<0.000001, 2-tailed Mann-Whitney test, **Fig 6-1G**) and were significantly less likely to select spontaneous routine turns (RT) than their siblings (dark: p<0.000001, light: p<0.000001, 2-tailed Mann-Whitney test, **Fig 6-1I**). Overall, mutants performed significantly fewer bouts per minute than their siblings in both dark and light conditions (dark: p<0.0001, light: p<0.0001, 2-tailed Mann-Whitney test, **Fig 6E**). Together, these results indicate that *ap2s1* regulates the baseline bout preference of larvae even in a spontaneous context.

### Neuronal expression of *ap2s1* is sufficient for acoustic habituation

Given the broad expression of *ap2s1* across neural and non-neural tissues ^50^, we sought to determine the cell types where *ap2s1* is important for behavioral regulation, focusing on acoustic response habituation, behavior selection, and kinematic performance. To determine if *ap2s1* regulates acoustically-evoked behavior through a direct neuronal function, we generated the *Tg(NBT:ap2s1-gfp)* transgenic line to pan-neuronally express GFP-tagged AP2σ in larvae. Transgene expression produced sustained green fluorescence broadly throughout the brain and spinal cord as early as 2 dpf **(Fig. 7A-B)**, allowing us to assess the behavioral impact of this neuronal tissue-specific expression.

**Figure 7.**
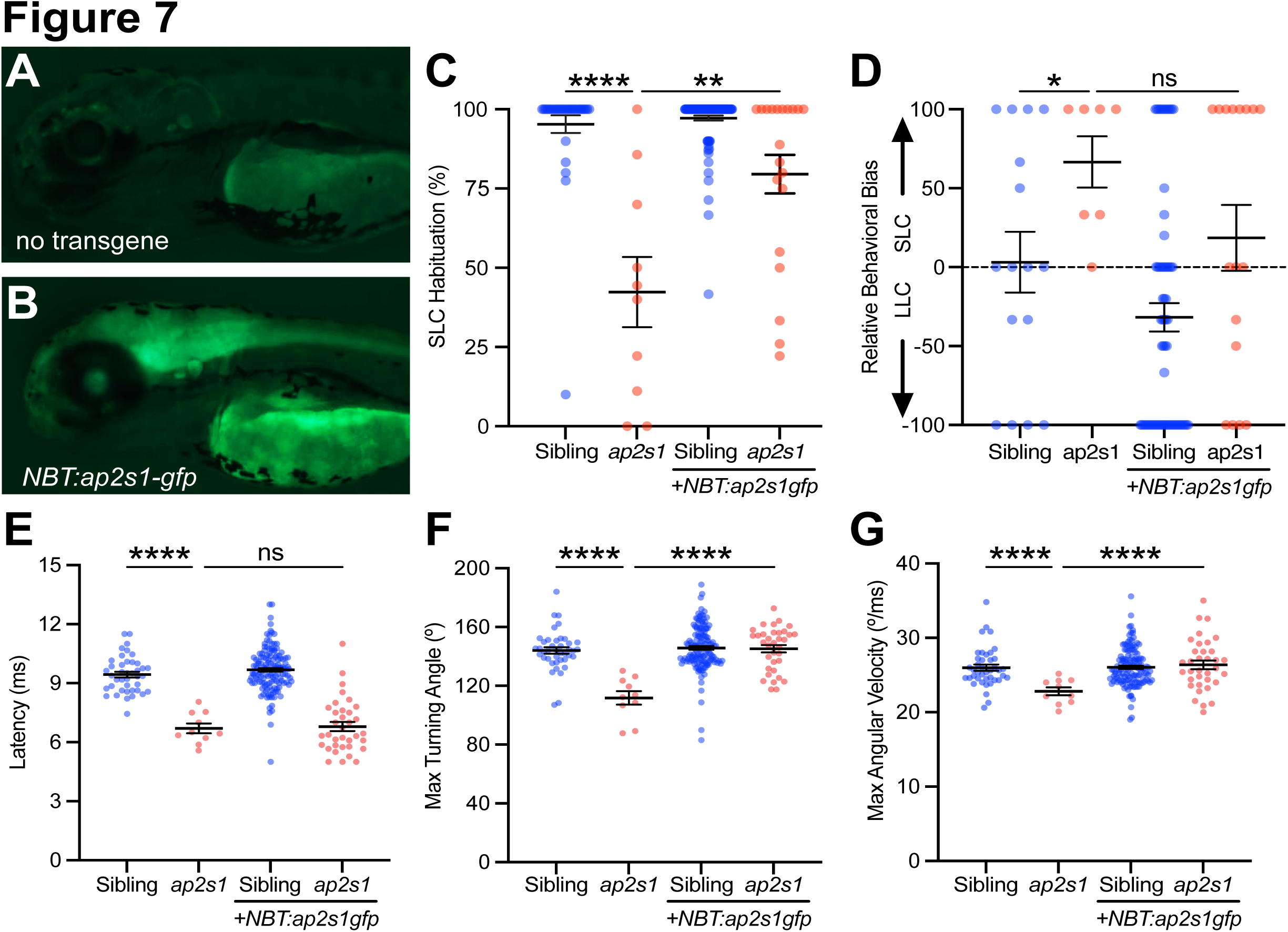
Neuronal expression of *ap2s1* is sufficient for acoustic habituation and response kinematics. **(A-B)** 3 dpf non-transgenic larva (A) and larva carrying *Tg(nbt:ap2s1-gfp)* (B) expressing green fluorescence in the brain and spinal cord. (**C**) Acoustic habituation percentage for sibling (blue) and *ap2s1^p172^* (red) larvae in the final 10 of 40 intense (23.8 dB) stimuli at 1s ISI (stimuli 31-40), with or without *Tg(nbt:ap2s1-gfp)* as indicated. **p<0.01, ****p<0.0001, Bonferroni corrected 1-tailed Mann-Whitney test. (**D**) Behavior selection bias of sibling (blue) and *ap2s1^p172^* (red) larvae in response to weak stimuli (-8.2 dB). *p<0.05, ns = not significant, Bonferroni corrected 1-tailed Mann-Whitney test. **(E-G)** Average kinematic parameters of SLC startle responses for sibling (blue) and *ap2s1^p172^* (red) larvae: latency (E), maximal turning angle (F), maximal angular velocity (G). ****p<0.0001, ns = not significant, 1-tailed t-test with Bonferroni correction. Mean and SEM indicated by black bars.

We first tested if neuronal *ap2s1* expression was sufficient to rescue the habituation deficits of *ap2s1* mutants. While acoustic habituation is compromised in non-transgenic *ap2s1^p172^* mutants compared to non-transgenic siblings (42.4% vs 95.3%, p<0.0001, Mann-Whitney test, **Fig 7C**), neuronal expression of *ap2s1-gfp* in *ap2s1^p172^* mutants significantly restored the ability to habituate (79.6%, p=0.0029, 1-tailed Mann-Whitney test, **Fig. 7C**). These data indicate that restoring neuronal expression of *ap2s1* is sufficient to modulate acoustic habituation. We next tested whether neuronal expression of *ap2s1* was also sufficient to regulate acoustically-evoked behavior selection. Compared to their siblings, *ap2s1^p172^* mutants show a significantly stronger bias towards selecting SLCs in response to low intensity stimuli (p=0.0282, Mann-Whitney test, **Fig 7D**). Expressing *Tg(NBT:ap2s1-gfp)* did not significantly rescue the response bias of *ap2s1^p172^* mutants back toward LLC’s, compared to non-transgenic mutants (p=0.1201, 1-tailed Mann-Whitney test, **Fig 7D)**, in contrast to the rescue observed for acoustic habituation. Finally, we examined whether neuronal *ap2s1* was sufficient to modulate the kinematic performance of acoustically-evoked responses, focusing on SLC latency, turning angle, and maximal angular velocity. Compared to their siblings, *ap2s1^p172^*mutant larvae initiate SLC bouts with a shorter latency (p<0.0001, 2-tailed t-test with Welch’s correction, **Fig 7E**), achieving a weaker SLC turning angle (p<0.0001, 2-tailed t-test with Welch’s correction, **Fig 7F**) and slower maximal angular velocity than their siblings (p<0.0001, 2-tailed t-test with Welch’s correction, **Fig 7G**). While neuronal expression of *Tg(NBT:ap2s1-gfp)* was not sufficient to restore the normal SLC latency in *ap2s1^p172^* mutants (p=0.397, 1-tailed t-test with Welch’s correction, **Fig 7E**), this neuronal *ap2s1* expression did significantly restore SLC turn angle (p<0.0001, 1-tailed t-test with Welch’s correction, **Fig 7F**) and maximal angular velocity (p<0.0001, 1-tailed t-test with Welch’s correction, **Fig 7G**), indicating that *ap2s1* regulates these kinematic aspects of acoustically-evoked behavior performance through neural-specific functions. Taken together, these data indicate that neuronal *ap2s1* activity is critical for regulating acoustic habituation learning and some response kinematics, though *ap2s1* likely modulates acoustically-evoked behavior selection bias and response latency through a different mechanism.

### Acute *ap2s1* activity is sufficient to regulate acoustic habituation

Since the AP2 complex has the potential to regulate neuronal function and behavior through developmental or acute activities, we tested the temporal requirements for *ap2s1* to understand the mechanisms through which this complex regulates acoustically-evoked behavior. We generated the *Tg(hsp70:ap2s1-gfp)* transgenic line to induce exogenous *ap2s1-gfp* expression in mutant larvae, using the heat-shock-inducible Hsp70 promoter for temporal control over AP2 complex activity ^69^. A single 38°C heat shock for 15 minutes induced ubiquitous green fluorescence, indicating robust *ap2s1-gfp* expression that persisted for >24 hours (**Fig 8A-C**). To determine whether the AP2 complex regulates acute circuit function to control acoustically-evoked behavior, we allowed mutant and sibling larvae to mature through 6 dpf, allowing development of the escape circuitry in the presence or absence of *ap2s1*, before inducing *ap2s1-gfp* expression via heat shock at 6 dpf. We then tested the acoustic habituation, behavior selection, and kinematic performance of larvae with and without the *Tg(hsp70:ap2s1-gfp)* transgene 12-18 hours later.

**Figure 8.**
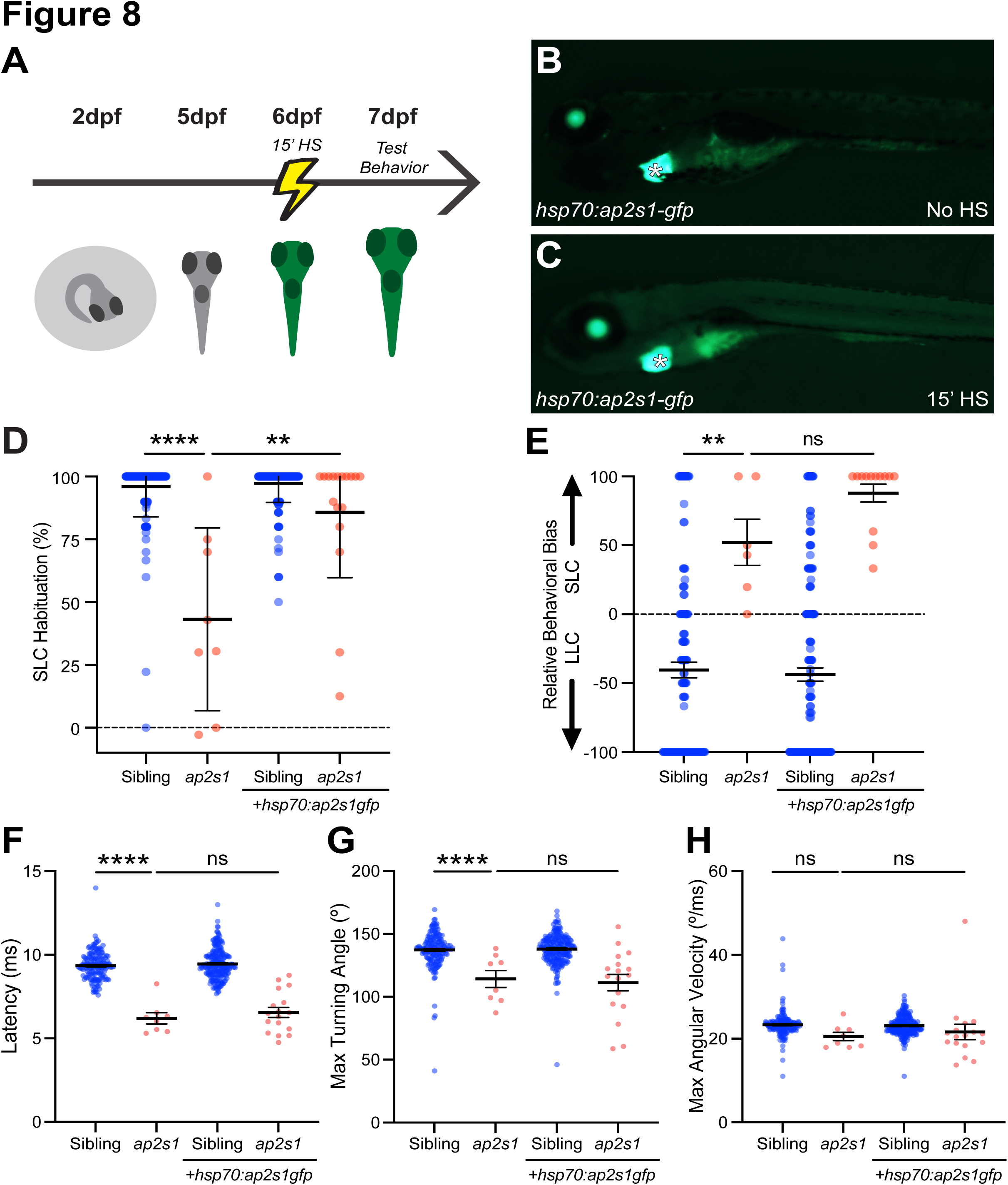
Acute expression of *ap2s1* is sufficient for acoustic habituation. **(A)** Timeline for acute induction of *ap2s1-gfp* expression in larvae. **(B-C)** 6 dpf larva carrying *Tg(hsp70:ap2s1-gfp)* without induction (B) and 5 hrs after a single 15 minute 38°C heat shock treatment (C). The transgene is marked with an independent GFP marker in the heart (asterisk). (**D**) Acoustic habituation percentage for sibling (blue) and *ap2s1^p172^* (red) larvae in the final 10 of 40 intense (23.8 dB) stimuli at 1s ISI (stimuli 31-40), with or without *Tg(hsp70:ap2s1-gfp)* as indicated. All larvae were treated identically with a heat shock. ****p<0.0001, **p<0.01, Bonferroni corrected 1-tailed Mann-Whitney test. (**E**) Behavior selection bias of sibling (blue) and *ap2s1^p172^* (red) larvae in response to weak stimuli (-8.2 dB). **p<0.01, ns = not significant, Bonferroni corrected 1-tailed Mann-Whitney test. **(F-H)** Average kinematic parameters of SLC startle responses for sibling (blue) and *ap2s1^p172^* (red) larvae: latency (F), maximal turning angle (G), and maximal angular velocity (H). ****p<0.0001, ns = not significant, Bonferroni-corrected 1-tailed t-test with Welch’s correction. Mean and SEM indicated by black bars.

Non-transgenic *ap2s1^p172^* mutants displayed habituation deficits relative to their non-mutant siblings following acute heat shock treatment (p<0.0001, 1-tailed Mann-Whitney test, **Fig 8D**). In larvae carrying the *Tg(hsp70:ap2s1-gfp)* transgene however, acute *ap2s1-gfp* induction at 6dpf significantly improved the acoustic habituation of *ap2s1^p172^* mutants relative to non-transgenic mutants (p=0.0058, 1-tailed Mann-Whitney test, **Fig. 8D**), demonstrating an acute impact of *ap2s1* on acoustic habituation. In contrast to the rescue of habituation, after acutely inducing *ap2s1-gfp* we did not observe any appreciable rescue of the acoustically-evoked behavior selection bias back toward LLCs (**Fig. 8E**). Finally, to determine if *ap2s1* regulates SLC kinematic performance through an acute mechanism, we tested for temporal-specific rescue of the *ap2s1*-dependent regulation of SLC initiation latency, turning angle, and maximal angular velocity. In the absence of *Tg(hsp70:ap2s1-gfp)*, heat shock treated *ap2s1* mutant larvae displayed shortened initiation latency, reduced SLC turning angle, and reduced maximal angular velocity (**Fig 8F-H**). Unlike for habituation, acute induction of *ap2s1-gfp* at 6dpf was not sufficient to appreciably rescue SLC latency (p=0.2274, 1-tailed t-test with Welch’s correction, **Fig 8F**), turn angle (p=0.3779, 1-tailed t-test with Welch’s correction, **Fig 8G**), or maximal angular velocity (p=0.3014, 1-tailed t-test with Welch’s correction, **Fig 8H**) in transgenic *ap2s1^p172^* mutants, suggesting that *ap2s1* regulates these kinematic aspects of acoustically-evoked behavior performance at an earlier timepoint, rather than through acute functions. Together, these data demonstrate that *ap2s1* acutely modulates acoustic habituation, while it may function through a distinct temporal mechanism to impact behavior selection and kinematic performance.

## DISCUSSION

Flexibility to select and perform contextually appropriate behaviors is critical for animals to survive and thrive. Here we demonstrate that the AP-2 complex broadly regulates behavioral choice and learning across environmental contexts. Human alleles of *AP2S1* produce complex behavioral effects ^20,25–27^, and while the AP2 complex plays highly conserved cell biological roles, connecting its cellular activity with resulting behavioral function has been challenging. *ap2s1* mutants are viable in zebrafish, unlike in rodents and *Drosophila* ^70,71^, providing the opportunity to directly probe the behavioral impacts of AP2 complex function through both neuron-autonomous and non-autonomous roles, distinguishing developmental and acute roles for AP2 in regulating behavior.

### The AP2 complex modulates multiple aspects of acoustically-evoked escape behavior

All three *ap2s1* alleles impact acoustic responsiveness, habituation, behavioral selection, and kinematic performance of escape behaviors (**Fig 1**), highlighting the varied critical roles for the AP2 complex in modulating when, where, and how escape behavior is carried out. Regulation of behavioral responsiveness, habituation, and selection are all conceptually similar, requiring the nervous system to process whether a given escape behavior occurs. However, genetic and molecular dissociation between these different modulations of escape behavior indicate that each is subject to separate regulatory mechanisms ^29,30,55,72,73^.

Increased acoustic responsiveness is frequently observed when acoustic habituation is impaired in zebrafish, both through diverse genetic disruptions including *pappaa*, *hip14*, *kcna1a*, and *cacna2d3* ^29,74,75^, and through pharmacological disruption of the NMDA-type glutamate receptor signaling with MK-801, ketamine, or 12-MDA ^55^. We demonstrate that *ap2s1* similarly coregulates acoustic responsiveness and habituation in zebrafish (**Fig 1**). This co-regulation appears broadly conserved as orthologs of *ap2s1* and many other ASD-associated genes regulate both responsiveness and habituation to mechanosensory stimuli in *C. elegans* ^33^. The AP2 complex also critically regulates selection between alternative escape behaviors, as both *ap2s1* and *ap2a1* mutations shift SLC vs LLC selection bias (**Fig 1 & 3**). Serotonergic and dopaminergic signaling appear critical for biasing this behavioral choice, suggesting possible neurotransmitter pathways that might be subject to AP2 regulation ^30^.

The shifts in movement kinematics of *ap2s1* and *ap2a1* mutants reveal the AP2 complex regulates behavioral performance of escape maneuvers in addition to behavior selection. The kinematic effects on SLC and LLC escape behavior performance dissociate (**Fig 2A-B, G-J**) and shift in opposing directions (**Fig 2M-O**), underlining the behavioral specificity of *ap2s1*’s functions. For example, while *ap2s1* mutants show a significant delay to initiate responses to dark flashes and shift to performing lower-energy turning behaviors (**Fig 5**), all *ap2s1* and *ap2a1* mutant lines bias their acoustically-evoked responses toward higher energy SLC behaviors (**Fig 1K-M**) that were consistently initiated with shorter latency (**Fig 2-3**). SLC latency shortens after ablating feedforward glycinergic inputs to the Mauthner neuron, pharmacological inhibition of glycine receptors, and *cacna2d3* mutation, which all also reduce SLC habituation ^75–78^. In contrast, the NMDA antagonist l-7,01,324 disrupts habituation through the feed-forward excitatory spiral fiber pathway with no impact on SLC latency ^79^, consistent with our transgenic dissociation of SLC habituation and latency. Together these suggest the AP2 complex could both enhance the activity of the feed-forward inhibitory glycinergic pathway to regulate acoustically-evoked SLC latency and regulate the feed-forward excitatory pathway to modulate habituation learning. Impaired feed-forward inhibition is a proposed mechanism underlying sensory processing and response shifts in individuals with autism ^80^, so detailing how *ap2s1* impacts acoustic feed-forward pathways will help model these effects in humans.

The AP2 complex uses different binding pockets and subunits to recognize and target a wide cohort of proteins for endocytic regulation, allowing for subunit specific roles ^81^. Indeed, single amino acid variants in human *AP2S1* can result in highly specific dysregulation of target proteins such as CaSR without disrupting the function of the overall AP2 complex ^54^. However, since *ap2s1* and *ap2a1* mutants show quite similar phenotypes in habituation, behavior selection, and kinematic performance, we argue that this reflects disrupting the overall AP2 tetramer complex rather than independent subunit roles. We do observe some phenotypic intensity differences across the *ap2s1* allele series. The extended habituation assay revealed that the *ap2s1^p172^* splice site mutation, which generates an in-frame protein deletion from the alternately spliced transcripts, has a milder effect on habituation rate and capacity than the *ap2s1^hv1^* deletion and *ap2s1^p199^* early truncation (**Fig 1**), suggesting the *ap2s1^p172^* protein product retains some function. An alternatively spliced *ap2s1* transcript producing an identical protein change to *ap2s1^p172^* has been detected endogenously in humans ^82^, suggesting the *ap2s1^p172^* splice variant may be functionally relevant. Milder substitution mutations of AP2 subunits, rather than deletions, may allow future dissection of the specific molecular targets regulating these diverse behavioral functions.

### *ap2s1* modulates acoustically-evoked behavior through multiple distinct circuit and temporal mechanisms

Our genetic rescue approaches demonstrate that even just in the context of a "simple" behavior such as the SLC escape, *ap2s1* modulates behavior through multiple mechanisms. By selectively restoring *ap2s1* function pan-neuronally through development (**Fig 7**), or acutely following escape circuit wiring and maturation (**Fig 8**), we demonstrate spatially and temporally distinct roles for the AP2 complex in learning, behavior selection, and behavioral performance. We propose there are at least three temporally and/or spatially distinct mechanisms through which *ap2s1* regulates acoustically-evoked escape behavior: 1) Acute Neuronal Activity, modulating acoustic habituation of SLC behavior, 2) Developmental Neuronal Activity, regulating SLC kinematic performance, and 3) Developmental Non-neuronal Activity, regulating escape behavior selection and response latency.

First, we propose that neuronal *ap2s1* functions after developmental wiring of escape circuitry to acutely modulate the rate and maximal acoustic habituation of SLC behavior. Pan-neuronally expressing *ap2s1-gfp* in *ap2s1* mutants was sufficient to rescue acoustic habituation learning (**Fig 7B**). The neuronal circuitry allowing acoustic habituation learning has already been established by 5 dpf in larvae ^55^, so inducing *ap2s1-gfp* expression at 5 dpf dissociates any potential developmental effects of *ap2s1* from acute roles in habituation. Since expressing *ap2s1* across 6-7 dpf is sufficient to rescue acoustic habituation, we conclude that *ap2s1* is critical for habituation after 5 dpf, and its activity is dispensable during circuit wiring for habituation. These results suggest the AP2 complex plays an active acute role in habituation learning, perhaps by promoting robust synaptic vesicle recycling to maintain behavioral inhibition or through postsynaptic receptor internalization to temporarily reduce excitatory signaling.

Second, we postulate that neuronal *ap2s1* functions prior to 6 dpf to regulate aspects of SLC kinematic performance, likely during the developmental wiring of the escape and/or motor circuitry. As we observed with habituation, pan-neuronal *ap2s1* expression was sufficient to rescue the weakened SLC turning angle and velocity of mutants. However, the *hsp70:ap2s1-gfp* experiments reveal a functional dissociation between habituation and the kinematic performance of escape behavior, as acute *ap2s1-gfp* expression significantly restored acoustic habituation learning without any corresponding effect on response latency or turning angle (**Fig 8D-G**). Since *ap2s1-gfp* was induced ubiquitously at 6 dpf, the critical window when *ap2s1* regulates SLC kinematic performance is at an earlier timepoint, likely during the developmental wiring of escape circuitry. While we cannot exclude the possibility that acute *ap2s1-gfp* expression at 6dpf was at an insufficient level to rescue response kinematics, this level of acute expression restored normal habituation in the same individuals with unrescued SLC kinematic performance, thus our experiments still demonstrate a clear dissociation between the functional levels and mechanisms of *ap2s1*’s regulation of SLC habituation and kinematics.

Third, we propose that *ap2s1* functions developmentally in a non-neuronal cell type to regulate acoustically-evoked behavior selection and response latency, as neither pan-neuronal, nor ubiquitous acute expression of *ap2s1* were sufficient to rescue escape behavior selection bias or appropriate SLC latency. For example, *ap2s1* is expressed in glial populations which could regulate formation of relevant synaptic connections ^31^, and also regulates PTH secretion and serum ion balance by endocrine cells ^83^, allowing non-autonomous effects on circuits through altered hormone or ion levels during development. While we cannot exclude the possibility that transgene expression levels were too low to rescue behavior selection or latency despite being in the right location or time, these same individuals demonstrated robust rescue of SLC habituation so the data nonetheless dissociate the mechanism of SLC habituation from behavior selection and SLC latency modulation. Examining additional candidate cell populations and inducing *ap2s1-gfp* expression during alternate developmental windows will be important to determine where and when *ap2s1* regulates escape behavior selection and SLC latency.

### AP2 broadly regulates behavior selection and learning across diverse contexts

By assessing behavioral responses across stimulus modalities and complexity, we demonstrate that *ap2s1* is a broad regulator of behavioral selection and learning. *ap2s1* modulates behavioral choice between distinct behavioral bouts in response to acoustic, visual, and mechanosensory stimuli, as well as in spontaneous contexts. In several contexts, *ap2s1* mutants shift their bias toward less vigorous movement bouts, suggesting that *ap2s1* promotes selection of higher intensity behaviors in these contexts (**Fig 5, 6**). Importantly, in acoustic contexts *ap2s1* mutants shifted toward the shorter latency and faster SLC responses, arguing against more trivial explanations of general sickness or low energy causing the observed behavioral changes. AP2 also regulates total bout performance frequency in a context-specific way: while AP2 has little effect on DF-evoked response rate, AP2 activity enhances spontaneous and loom-evoked bout frequency, while reducing acoustically-evoked response frequency. Visually evoked behaviors of DF, OMR, and prey capture assays are independent of the escape response circuitry ^63,84^, thus *ap2s1* likely regulates these behaviors through distinct circuit mechanisms. Weak spontaneous turns and stronger visually-evoked turns are thought to be controlled by a shared subset of spinal projection neurons of the brainstem whose activity is critical for selecting turn vs swim behaviors ^65^, making these neurons candidate loci for *ap2s1*’s regulation of visual and spontaneous behavior selection.

The modality-specific impacts of *ap2s1* on escape behavior selection highlight multiple critical roles for the AP2 complex in information processing in the brain at diverse circuit locations. Although acoustic, visual, and mechanosensory stimuli all engage the SLC and LLC escape circuitries, our data suggest that AP2 function promotes LLC selection in response to acoustic stimuli while promoting SLC selection in response to visual and mechanosensory modalities. Mechanistically, this suggests that *ap2s1* regulates escape behavior selection at the level of information processing and/or integration of sensory information, rather than simply regulating the excitability of shared command and motor neurons involved in escape behavior execution. Determining how *ap2s1*’s temporal and spatial requirements for visual or mechanosensory escape behaviors compare to those of acoustic escape behaviors will shed light on whether *ap2s1* has independent roles in modality-specific processing or a more centralized role in multisensory integration.

Habituation reflects experience-driven shifts in behavior selection biases and we show the AP2 complex regulates this as well. Disrupting *ap2s1* results in a dramatic deficit in habituation to repeated acoustic stimuli (**Fig 1**), though disrupting *ap2s1* enhances the habituation of visually-evoked turns (**Fig 5G**), suggesting AP2 plays multiple diverse roles in short-term habituation across sensory modalities. DF habituation is mechanistically more complex than SLC habituation, also involving kinematic weakening and a progressive delay in response latency through parallel molecular pathways ^64^. Since *ap2s1* mutants already show a significant shift in DF response latency under non-habituating conditions, possible roles for *ap2s1* in regulating these other aspects of visual habituation remain unclear. Surprisingly, the OMR directionality of *ap2s1* mutants was also significantly enhanced over their siblings during the optomotor assays (**Fig 6A-B**). We speculate *ap2s1* could regulate habituation to these visual stimuli and mutants then fail to reduce their responses across the assay, though more extended OMR assays with finer temporal resolution would be required to test this possibility ^85^. Alternatively, biasing bout selection toward less intense bouts as observed during spontaneous swimming (**Fig 6D**) might help mutants maintain the correct heading.

In humans, function-disrupting *AP2S1* alleles can result in learning disabilities, cognitive deficits, and/or behavioral disturbances. These alleles have been linked to ASD and ADHD in children, though the specific behavioral changes associated with these *AP2S1* cohorts are just beginning to be elucidated ^25,26,28^. The diverse behavioral impacts of *ap2s1* disruption in zebrafish echo many behavioral phenotypes of adolescents with ASD. Increased sensitivity and enhanced behavioral responses to acoustic or mechanosensory stimuli are some of the most common sensory phenotypes of ASD ^80,86–88^ and children with ASD show reduced habituation of behavioral responses to repeated sounds ^89,90^. Visual sensitivity and visually-evoked responses are often altered in populations with ASD ^91,92^, where some repeated visual stimuli evoke less neural activity habituation ^93^. Notably, observed behavioral shifts vary considerably across cohorts and populations with ASD, reflecting the genetic diversity underlying the autism spectrum ^80^. Thus, the zebrafish *ap2s1* model suggests behavioral phenotypes to explore specifically in *AP2S1*-linked ASD, helping to illuminate the breadth of its conserved roles in more complex learning and behavior selection.

## METHODS

### Zebrafish Maintenance

All fish were maintained on a 14h:10h light:dark cycle. Zebrafish larvae and embryos were raised and maintained in 1X E3 solution at 29°C unless otherwise noted. All mutant and transgenic lines were maintained in the Tüpfel Long Fin (TLF) strain (Jain et al., 2018). Sex is not determined in zebrafish until 25-60 dpf so behavioral analyses of larvae were performed without consideration of sex. All experiments with zebrafish (*Danio rerio*) were approved by the Haverford College IACUC and/or the French Ministère of Research and the Ethic Committee of the Sorbonne Université and Institut Curie. Animal handling and experimental procedures conducted at the Champalimaud Centre were approved by the Champalimaud Centre for the Unknown Bioethics Committee and the Portuguese Veterinary General Board (Direcção Geral Veterinaria), in accordance with the European Union Directive 2010/63/EU.

### Generation of Mutants and Transgenic lines

The novel *ap2s1^hv1^* mutation was generated using CRISPR as previously described for *ap2s1^p199^* ^30^, using a sgRNA targeting the genomic sequence 5’-GGAAGACCAGACTGGCCAAGTGG-3’. The *ap2a1^sa1907^* allele was a TILLING mutant obtained from ZIRC. To generate the transgenic constructs, full length wild type *ap2s1* cDNA was amplified from TLF zebrafish larvae into pENTR/D-TOPO (ThermoFisher), inserting an in-frame 15bp flexible linker sequence (encoding the amino acids GGGGA) followed by EGFP at the C-terminus using Infusion cloning (Clontech). For the *Tg(NBT*:*ap2s1-gfp)* construct, *ap2s1-gfp* was cloned into a destination vector under control of the *Xenopus* NBT promoter for pan-neuronal expression ^69^ and followed by the SV40 polyadenylation sequence. The full transgene was flanked by I-SceI meganuclease cleavage sites, and transgenic lines were generated by coinjection with I-SceI (NEB) as previously described ^94^. For the *Tg(hsp70*:*ap2s1-gfp)* construct, *ap2s1-gfp* was cloned into the pDestTol2CG2 vector between the heat-activated *hsp70* promoter to drive inducible *ap2s1-gfp* expression in all tissues and the SV40 polyadenylation sequence. This transgene also carried an independent *myl7:gfp* marker expressing GFP in the heart, and we generated transgenic lines by coinjection with *tol2* transposase mRNA ^95^.

### DNA Preparation and Genotyping

Mutant *ap2s1* and *ap2a1* alleles were detected as previously described (Jain et al., 2018), where larval genomic DNA was extracted ^96^, then PCR amplified with GoTaq MasterMix (Promega) and subsequently digested with a restriction enzyme to distinguish wild type and mutant alleles (**Table 1**). PCR reactions were supplemented with 5% DMSO for *ap2s1^p172^. ap2s1^p172^* alleles were distinguished by gain of a BsmAI (NEB) cutting site, *ap2s1^p199^* and *ap2s1^hv1^* alleles were distinguished by loss of a MscI (NEB) cutting site, and *ap2a1^sa1907^* alleles were distinguished by gain of a HinfI (NEB) cutting site. PCR amplification used Tanneal 48°C for *ap2s1^p172^* and 54°C for *ap2s1^p199^* and *ap2s1^hv1^* with the following program: 2 min - 94°C, 40×[30s - 94°C, 30s - Tanneal, 45s - 72°C],10 min - 72°C. Amplification of *ap2a1^sa1907^* used the following program: 2 min - 94°C, 35×[30s - 94°C, 30s - 55°C, 45s - 72°C], 10 min - 72°C. Transgenic larvae were molecularly verified by PCR using primers to detect *gfp* sequence (Table 1) and the following program: 2 min - 94°C, 30×[30s - 94°C, 60s - 59°C, 60s - 72°C], 5 min - 72°C.

**Table 1:**
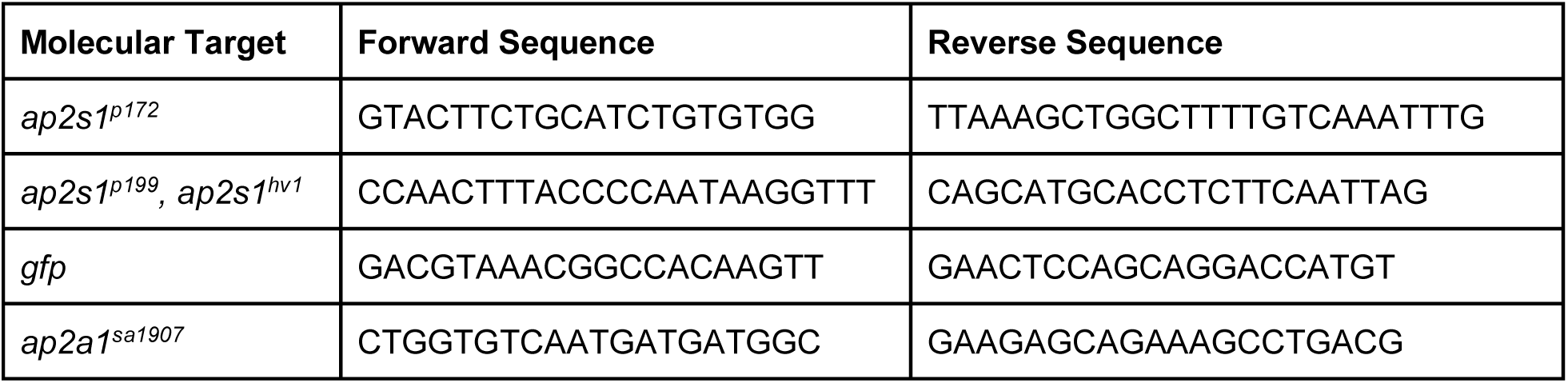
Primer sequences used for genotyping

### Behavioral Testing and Analysis

All larval behavior was assessed at 5-7 dpf, and any larvae with gross morphological abnormalities at testing time were excluded. Since *ap2s1* mutants sometimes show variable delays in swim bladder inflation, fish with and without swim bladders were both included in experiments to avoid selecting against mutants. All behavioral experiments were carried out blind to genotype with subsequent genotyping of all individuals, always comparing mutants to their siblings tested and handled in the same experiment.

#### Acoustically-Evoked Behavior

After acclimating to testing conditions on a light box for 45min, healthy fish were placed in individual wells of a custom plexiglass testing arena, either 9mm diameter wells in a 36-well arena or 9mm × 9mm square wells of a 16-well arena, illuminated from below with an IR array (CM-IR200-850, CMVision), attached to a vibrational exciter (4810, Brüel & Kjær) that administers acoustic stimuli, as previously described ^30,97^. Acoustic stimuli were 2ms tones (1000 Hz), varying intensity as needed for each assay. Stimuli were generated and triggered using DAQTimer software ^63^, and videos were recorded in 120 ms bursts initiating 30 ms prior to each acoustic stimulus with a high-speed camera (TS4, Fastec) at 1000 fps. To test larvae for bias and sensitivity phenotypes, acoustic stimuli of varying intensities (-8.2 dB to 25.9 dB) were delivered interleaved at 20 sec ISI for 10 stimuli at each intensity. Larvae were tested for acoustic habituation by presenting 10 pre-habituation stimuli at 20s ISI (23.8 dB) followed by 30 or 300 more identical stimuli at 1sec ISI. Video files were tracked and analyzed using FLOTE ^63^, using a latency threshold of 15 ms in separating SLC from LLC behavior. Larvae responding to fewer than 40% of pre-habituation stimuli were excluded from habituation analysis. SLC Habituation scores for the larvae were calculated in blocks of 10 stimuli [SLC habituation score = 100*(current block – pre-habituation)/pre-habituation]. For the extended habituation assay, a least squares fit hyperbola was generated for each data set using Graphpad Prism 9, with "Bmax" representing the maximal habituation asymptote and "Kd" representing the half-maximal habituation.

#### Recording of mechanosensory, chasing dot, and spontaneous behavior

The behavioral setups used for the mechanosensory startle assay, chasing dot assay, and spontaneous swimming in the light and dark have been previously described ^53^. The fish were recorded from above at 700 frames per second (shutter time 1423 µs) using a high-speed infrared sensitive camera (MC1362, Mikrotron-GmbH, Germany) equipped with a Schneider apo-Xenoplan 2.0/35 lens. The larvae were illuminated from bellow with a 10 × 10 cm LED-based diffusive backlight (850 nm) and the camera was fitted with a 780 nm long pass filter that blocked visible light. Fish tracking and segmentation of the tail were performed online using a custom-made algorithm in C#, as previously described ^98^.

#### Mechanosensory startle assay and spontaneous (dark) behavior

The mechanosensory startle assay was adapted from Groneberg et al ^59^. A 2 cm long capillary rod, made of the tip of a loading-pipette (outer diameter: 0.3 mm), was attached to a piezoelectric bender actuator (Thorlabs; voltage range: 150 V; maximum displacement: 450 µm). The capillary rod was immersed 2 mm into the arena’s water and mounted at a 45° angle. Single fish were placed in circular acrylic arenas with 2.5 cm of diameter and 3 mm depth for 1h30min. The piezo deflections were composed of five 2 ms pulses at a fixed 80 V voltage with a 2 ms inter-pulse interval. To avoid habituation, stimuli were triggered with a minimum inter-stimulus interval of 2 min. To ensure that the larva was always facing away from the rod the piezo was triggered only when the larva was at a distance of 4 mm of the rod’s tip and swam to at least a 5 mm distance from it. After, there was a 400 ms delay to activate the piezo to guarantee that the fish was not performing a bout when the stimulus was triggered. To ensure that fish swam to the vicinity of the capillary rod and were able to activate it, circular sine wave converging gratings (spatial period of 10 mm, velocity 5 mm/s) centered at the tip of the piezo were presented underneath the fish by a DLP projector (BenQ). Mechanosensory-induced startle bias was calculated using the formula: 100% × (SLC frequency – LLC frequency)/(total SLC + LLC response frequency), where score of 100% results from only SLCs and a score of -100% from LLCs. The spontaneous swimming in the dark assay was performed in the same behavioral set up and arenas as the mechanosensory startle assay, but the projector was turned off ensuring that the fish was in total darkness (0 lux) and the epochs where the fish activated the piezo were excluded from the analysis.

#### Chasing dot and Spontaneous (Light) Behavior

Single larvae were placed in an acrylic circular arena with 5 cm of diameter and 4.5 mm depth. The arena is completely transparent so that visual stimuli could be displayed even when the larva was swimming in its borders and had a round bottom as animals spend less time in its borders compared with arenas with a flat bottom and vertical edges. The visual stimuli were projected by an Optoma ML 750e projector onto a diffusing screen placed 1.5mm below the larva. In the spontaneous swimming assay a homogenous white square with 1000 lux was displayed for 20 min. In the chasing dot assay, a dark spot of 1 mm radius was projected 2 cm away from the larva and approached it with a speed of 0.5 cm/s and was updated in real time to stay locked to the fish’s position and orientation. When the dot reached the larva, it stayed underneath it for 1 s. The dot approached the larva from the left or the right, and both directions of stimuli were presented with an inter-stimulus interval of 5 min. The background during stimulus and inter-stimulus periods was the same homogenous white square displayed for the spontaneous swimming in the light assay. Chasing dot stimuli were presented 7 times from each direction per animal, 14 stimuli total.

#### Visual Loom Behavior

6dpf larvae were adapted to the behavior arena conditions for at least 2 hours prior to the experiment, and individually tested in 35mm petri dishes in E3 media. Video was acquired at 20.2 pixels/mm at 750 Hz using a high-speed camera (MC4082, Mikrotron-GmbH, Germany) and a Kowa LM35HC 1" 35mm F1.4 lens, using an Active Silicon FireBird frame grabber board (P/N: MP-FBD-4XCXP6-2PE8). The behavioral arena was illuminated from below with an infrared LED array, and an 850 nm infrared bandpass filter (BP850-35.5, Midwest Optical Systems, Inc.) was used on the lens to block all the visible light. A custom-written C# software extracted the fish position and orientation, and updated stimulus position in real-time in a closed loop (adapted from Dunn et al ^60^). Stimuli were projected on a plexiglass screen underneath the fish using a cold mirror and a DLP projector (Optoma). Stimuli were presented to either the left or right side of the fish, centered 0.5 cm from the center of mass of the fish. The stimulus expanded with a velocity of 250 mm/s and disappeared 6 s after the expansion began or after a high-velocity escape response (calculated based on a moving average) was performed by the larva. The analysis of the escape response induced by the looming stimulus was carried out using a custom-written MATLAB script, available on request.

#### Optomotor Behavior

6dpf larvae were individually tested in a 9cm clear petri dish resting on a white plexiglass sheet, using the same imaging system as Visual Loom experiments. Larvae were presented with a black/white grating of 5 mm period projected from below. The gratings moved perpendicular to the stripe direction at 5 mm/s for 30s, followed by 10s of the non-moving grating. Stimuli were presented 10x, alternating between left-and rightward stimulus movement, and larval behavior was captured at 20 Hz from above. Head position was tracked offline using DeepLabCut ^99^. Analysis pipeline and path plotting was done with a custom MATLAB script, available on request. The OMR score was calculated with the formula: 100*(distance traveled in stimulus direction – distance traveled opposite stimulus direction)/total distance traveled. Only coordinates within a cropped region of the arena were included for directionality and path plotting analysis, to exclude data points where fish encountered the edge of the dish. Fish paths that reentered the cropped arena during the trial were not depicted in **Fig 6B** for clarity, though they were included in OMR score. Velocities were smoothened by excluding data points exceeding a 9 pixel/frame threshold.

#### Dark Flash Behavior

Larvae were acclimated to constant illumination for at least 3 hours prior to testing, then transferred to the array of 36 individual 9mm diameter testing arenas used for acoustically-evoked behavior, lit obliquely above with a dimmable white LED (MCWHL5, Thor Labs). Behavior was recorded at 500 fps, triggered to start recording 30ms before each "Dark Flash" where the white LED was turned off for 2 seconds. All components were surrounded by a custom black vinyl enclosure to shield from external lights. 6dpf larvae were presented with 6x “pre-habituation” dark flashes at 150s ISI, followed immediately by 42 “habituating” dark flashes at 15s ISI. Testing took place between the hours of 10:00am and 5:00pm. Videos were tracked and analyzed for response rates and response latency using Flote v2.1 ^63^, and subsequent bout classification was performed through an offline bout-based tracking approach (see *“Bout-based Tracking”* section below).

#### Rotifer Consumption

Imaging set-up was the same as employed in the looming and OMR experiments but without the infrared bandpass filter as a dark field illumination was created using a low-angle white-light LED ring (CCS, Japan; model: LDR2-100SW2-LA) to allow visualization of both the larva and rotifers. Image acquisition frequency was 20 Hz in 10s bursts. Sets of 60-80 6 dpf larvae were pre-fed with >1000 rotifers in 90mm petri dishes for 3 hours, then rotifers were washed out with fresh filtered E3. Individual larvae were transferred to 60mm dishes containing 5ml of 5g/L sea salt (Instant Ocean Salt) and starved for 24 hours. Starved 7dpf larvae were given approximately 100-120 rotifers per fish per dish to hunt. Starting immediately upon addition of rotifers (t= 0 min), 10s bursts were recorded from each dish at five time points across 135 minutes of the experiment. Each individual larva was saved for subsequent genotyping. Videos were analyzed using “Find Maxima” in FIJI ^100^, counting the total number of visible rotifers in each frame of each video. Since some rotifers could move in and out of video detection at the edge of the chamber, the frame with the largest number of particles was used to measure a single rotifer count for each time point from, subtracting the number of detected particles that represented parts of the fish larva rather than rotifers. Rotifer counts were normalized to 100% against the initial prey count at t= 0 min for each fish to assess rotifers consumed by each larva across the assay.

#### Bout-based Tracking

The swim bout type classification has been described in detail in ^53^. Bouts and tail half-beats were detected and this information was used to compute, for each swim bout, 73 kinematic parameters as described previously ^53^. This information was used to embed the data into a previously computed principal component analysis (PCA) space that was obtained from a data set of 3 million bouts that fish executed over a wide range of behaviors and stimuli ^53^. To categorize the bouts into types, we used a dataset composed of equal number examples of each bout type (Bout Map) ^53^, that were categorized using density valley clustering (clusterdv)^101^, and used k nearest neighbors (k = 50) over the first 20 principal components to assign every new bout to one of the possible 13 categories. To apply the same analysis pipeline to the dark flash data acquired using a different set-up and frame rate, an offline version of the tracking algorithm from Marques et al.^53^ was applied to the recorded movies, and the data was interpolated to 700 frames per second. To visually compare the kinematic distributions in mutant and sibling groups, scatter plots were made using the first two PCA dimensions from Marques et al. 2018^53^. Changes in the distribution of kinematics between groups were visualized by taking the difference between kernel density estimates based on 2D Gaussian kernels (Matlab ksdensity function) in the first two PCA dimensions.

### Transgenic Rescue

Pan-neuronal rescue of *ap2s1* was carried out using an incross of *Tg(NBT:ap2s1-gfp); ap2s1^p172^*/+ adult zebrafish. Larvae were genotyped for both *ap2s1* and the transgene after behavioral testing. Acute rescue of *ap2s1* was carried out using an incross of *Tg(hsp70:ap2s1-gfp); ap2s1^p172^*/+ adult zebrafish. Larvae were pre-screened for transgene presence through heart fluorescence from the *myl7:GFP* marker on the transgene, and *ap2s1* genotype was molecularly determined after behavior testing. The *hsp70* promoter is ubiquitously inducible by abrupt heat shock, though as previously described the promoter also drives targeted expression in the lens without heat shock induction ^102^. To induce *ap2s1-gfp* expression, unanesthetized larvae were transferred to individual wells of a PCR plate in 100µl E3 to receive a 38°C heat shock for 15 min. Following heat shock, larvae were returned to 6cm dishes in fresh E3 at 29°C. Transgenic fish were always compared with non-transgenic siblings that also received the same heat shock regimen to control for background and handling.

### Statistical Analyses

2-tailed Mann-Whitney tests were used to compare end-point habituation, bias, and responsiveness scores, where floor or ceiling values produced non-Gaussian distributions (GraphPad Prism 9). 2-tailed unpaired t-tests with Welch’s correction for unequal variance were used to compare kinematic parameters (GraphPad Prism 9). For experiments where bout types were categorized (mechanosensory startle assay, chasing dot assay, spontaneous swimming in the light and dark) we applied a two-tailed Wilcoxon signed-rank test (MATLAB: ranksum), and adjusted p-values were corrected for multiple comparisons using the Holm-Bonferroni method. Two-way repeated measures ANOVA analyses were used to assess the significance of differences across time and genotype for response likelihood measures of the acoustic and DF habituation assays. R Studio was used to compute Hartigan’s dip test for multimodality on Dark Flash latency distributions. 1-tailed statistical tests were used in assessing transgenic rescue, as these tested for a phenotypic shift in a single direction. The Bonferroni correction was used to conservatively account for multiple comparisons with single datasets. Unless otherwise indicated, graphs display mean ± SEM.

## Acknowledgements

Funding: J.C.M. was funded by H2020-MSCA-IF-2019 (897403). G.R. was supported by a European Union’s Horizon 2020 research and innovation programme under the Marie Skłodowska-Curie grant agreement No 666003. M.B.O. and A.L. were supported by FCT (Portugal; PTDC/MED-NEU/32664/2017), Volkswagen Stiftung “Life?” Initiative and ERC (NEUROFISH 773012). R.A.J. was funded by the National Eye Institute of the NIH (R15EY031539). We thank Clarice Xu, Emilia Cobbs, Christina Szi, and Karine Durore for experimental assistance, and Dr. Eric Miller & Dr. Jessica Nelson for discussion and manuscript feedback. We thank the Champalimaud Molecular and Transgenic Tools Platform (MTTP) and Fish Facility (supported by *Congento LISBOA-01-0145-FEDER-022170, co-financed by FCT (Portugal) and Lisboa2020, under the PORTUGAL2020 agreement (European Regional Development Fund)* for logistic support.

---

**Supplemental Figure S1:**
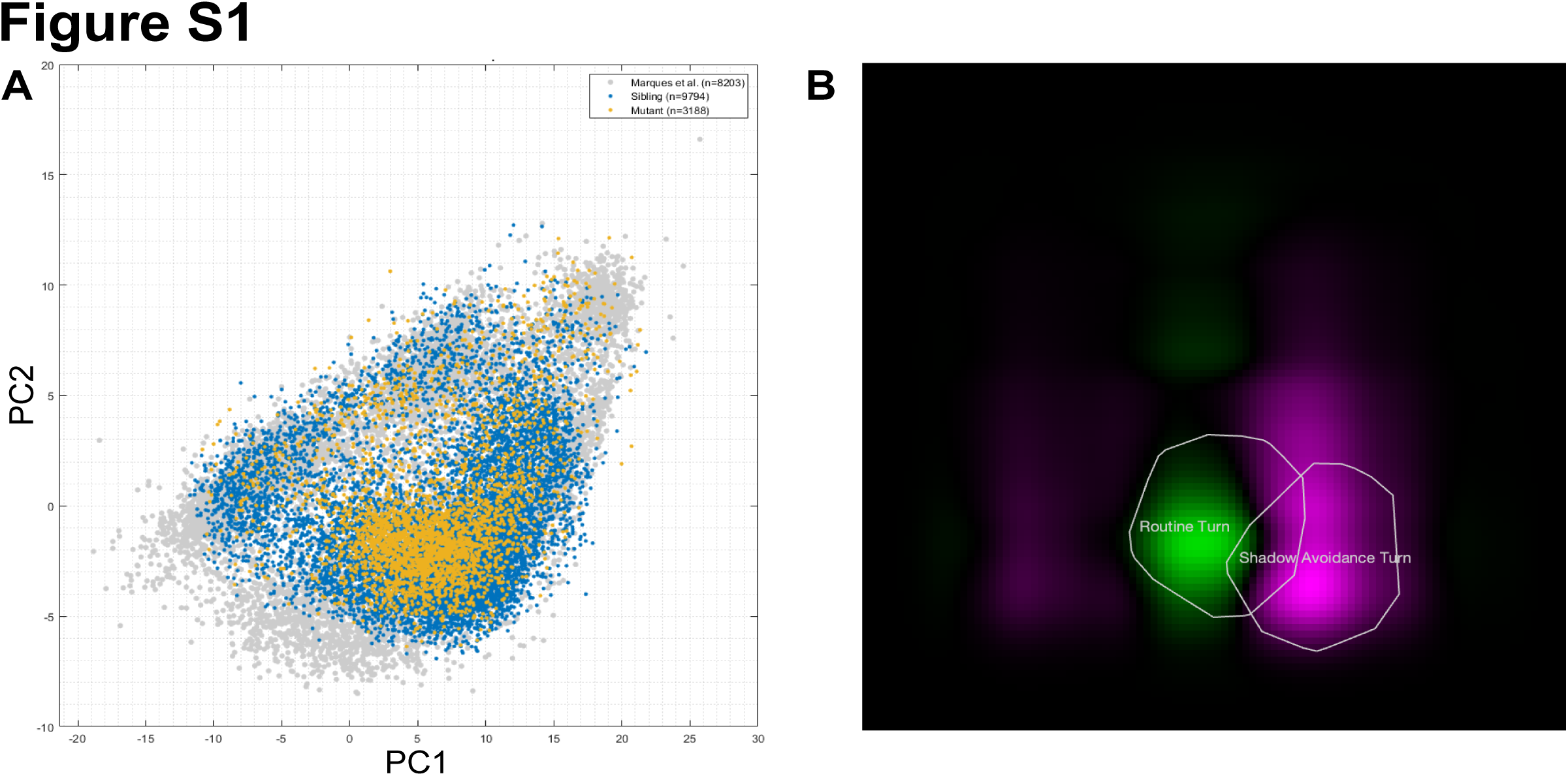
Spontaneous behavioral bout selection by *ap2s1^p172^* mutants and their siblings. **(A)** Distribution of DF-induced bouts in kinematic PCA space from *ap2s1^p172^* mutants (yellow) superimposed on non-mutant siblings (blue) and bouts from Marques et al. 2018 (grey). All data sets have similar extent, although Marques et. al data set also contains bouts relating to prey capture (bottom left) and fast escapes (top right) not observed in our spontaneous recordings. (**B**) Difference in bout density in kinematic PCA space between *ap2s1^p172^* mutants and non-mutant siblings. Green indicates an increase in bout probability in mutants, and magenta a decrease. Each color is normalized to the maximum of the distribution. White rings show the typical distribution of RTs and SATs, in the form of the convex hull surrounding 90% of bouts closed to the cluster center.

**Supplemental Figure S2:**
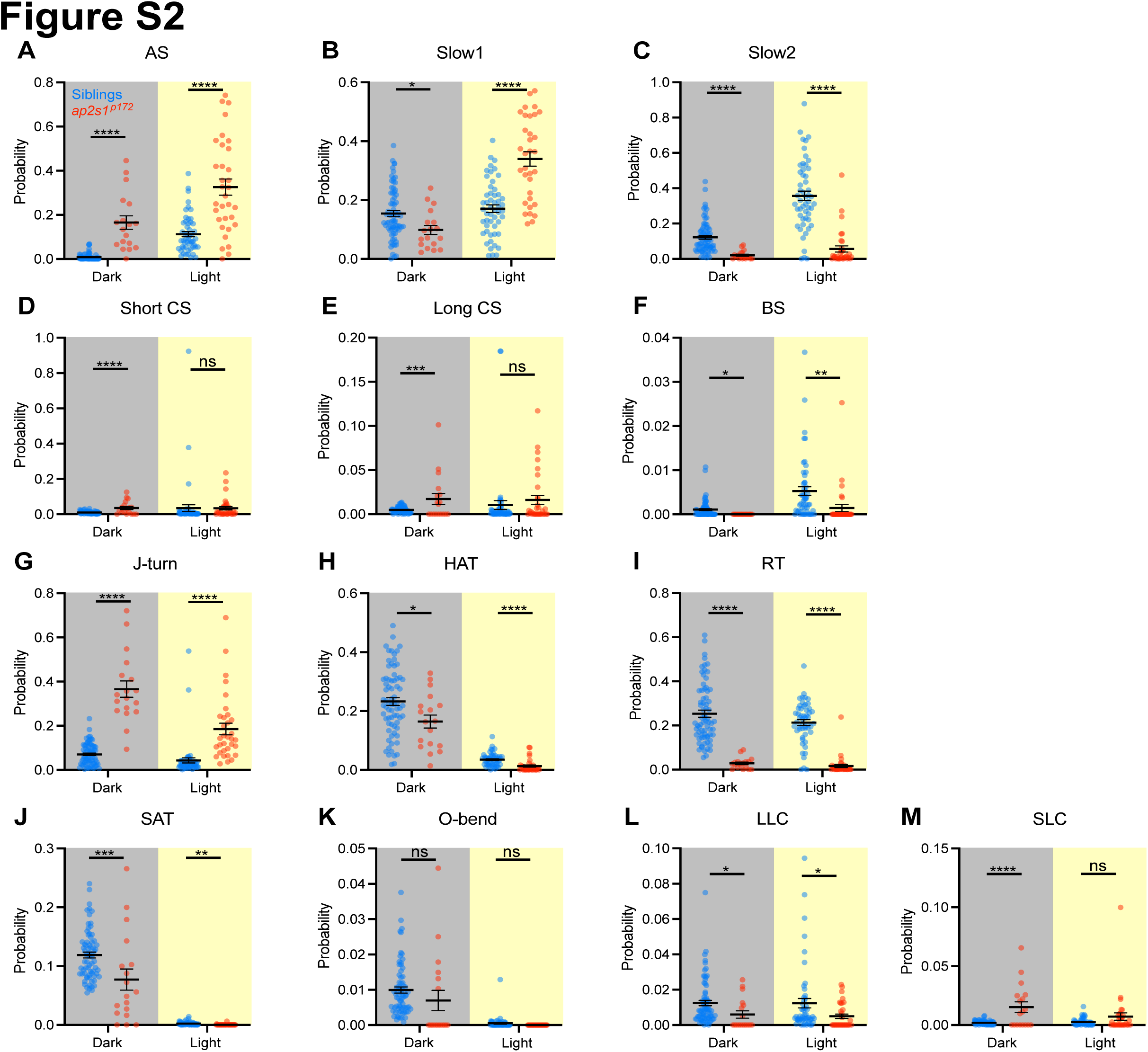
Spontaneous behavioral bout selection by *ap2s1^p172^* mutants and their siblings. Frequency of bout selection over 20 minutes of spontaneous activity in uniform darkness or light by individual non-mutant siblings (blue, n=72 in dark, n=52 in light) and *ap2s1^p172^* mutants (red, n=18 in dark, n=33 in light). **(A-M)** Behavioral bouts are classified as in Marques et al. (2018), including Approach Swim (AS), Slow Swim 1 (Slow 1), Slow Swim 2 (Slow 2), Short Capture Swim (Short CS), Long Capture Swim (Long CS), Burst Swim (BS), J-turn (J-turn), High Angle Turn (HAT), Routine Turn (RT), Spot Avoidance Turn (SAT), O-Bend (O-Bend), Long Latency C-start (LLC), and Short Latency C-start (SLC). Significance was assessed through 2-tailed Mann-Whitney test with Bonferroni correction: *p<0.00385, **p<0.000769, ***p<0.0000769, ****p<0.00000769. All error bars indicate SEM.

